# Electron microscopy reveals water networks inside hydrated cellulose fibers

**DOI:** 10.64898/2026.09.16.752152

**Authors:** Ruoya Ho, Arunabh Athreya, Yanina Pankratova, Chaemyeong Lim, Louis Wilson, Mei Hong, Keisuke Nakashima, Jochen Zimmer

## Abstract

Cellulose is a highly abundant linear glucose polymer that exists in amorphous and fibrillar forms^1-3^. Primarily produced by vascular plants but also microbes and tunicates, cellulose predominantly performs architectural functions by stabilizing cell walls, tissues, and multicellular communities^4-9^. Owing to its association with other cell surface materials, physiological higher-order structures of cellulose have been difficult to obtain. Here, we describe a never-dried tunicate cellulose fibril structure by cryogenic electron microscopy, which resolves a fiber of more than 360 cellulose strands. Our data reveals linear and severely twisted fiber segments that are interspersed with solvent channels. Solid-state NMR analyses and molecular dynamics simulations indicate the presence of structural water molecules that form a network connecting neighboring cellulose chains. Confocal and super-resolution MINFLUX fluorescence imaging using cellulose-specific probes as well as biochemical analyses demonstrate shared surface and material properties of plant and tunicate cellulose fibers. Combined, our data suggest that architectural water molecules may be common features of cellulose fibers and likely other polysaccharide assemblies in their hydrated physiological states.

## Main

Cell walls are essential for terrestrial plant growth and contain load-bearing cellulose fibrils that encircle the cell ^1^. First described by Anselme Payen in 1838 ^2^, cellulose is a linear β-(1,4)-linked glucose polymer that is a versatile building block for many biomaterials ^3^. Cellulose’s biophysical and material properties have been characterized in detail, including its organization in plant cell walls, bacterial biofilms, and animal tissues ^4-9^. In tunicates, cellulose is a structural component of the tunic, an integumentary tissue covering the entire animal ^10-13^.

Cellulose is synthesized and secreted by cellulose synthase ^14^, and multiple polysaccharides most likely assemble spontaneously into fibrils under suitable conditions ^15^. Cellulose fibers are produced by plants, certain bacteria and fungi, as well as tunicates. The fiber architecture is likely important for its material properties, including interactions with other cell wall components, such as hemicelluloses and lignin ^16-18^. Diffraction studies of dried cellulose crystallites yielded high-resolution insights into the crystalline forms of algal and tunicate cellulose fibers, defining the cellulose I_α_ and I_β_ allomorphs, respectively ^19,20^. Importantly, these seminal contributions established a rigid cellulose fibril architecture devoid of solvent.

Detailed high-resolution structural analyses of native hydrated cellulose fibers have been challenging due to their association with other matrix biopolymers in plant tissues. Here, we employed cryogenic electron microscopy (cryo-EM) to determine the structure of a native-like, never-dried pure cellulose fibril from the tunicate C*iona intestinalis* ^*12*^. Our analyses reveal a rhomboidal fibril architecture of cellulose strands separated by water channels. Within a layer, neighboring cellulose chains interact via hydrogen (H)-bonding to structural water molecules, and layer stacking is accomplished by van der Waals interactions. Solid-state NMR analyses and molecular dynamics simulations support the presence of structural water molecules throughout the fibril at the interface of four cellulose strands. Using confocal and super-resolution fluorescence microscopy, we demonstrate that the hydrated tunicate fiber shares many properties with plant cellulose fibrils, suggesting that the observed structure represents fibrillar cellulose in its physiological hydrated form.

### Tunicate cellulose fibers are long and flexible

Never-dried cellulose fibrils from the tunicate *Ciona intestinalis* were harvested and TEMPO oxidized to increase water dispersibility (see Methods) ^21^. Imaging by negative-stain electron microscopy revealed well-separated fibers that occasionally formed bundles and overlapped with other fibrils (Fig. 1a). Images capturing the ends of a continuous fibril suggest a length of at least 7 μm (Fig. 1a). Under cryogenic conditions, we also observed separated but occasionally aggregated fibers in different orientations (Extended Data Fig. 1). The fibers exhibit thicker and thinner segments, like a randomly twisted ribbon. For structural analysis, we employed a single-particle cryo-EM processing approach by picking particles of 350 Å diameter along the fibrils (Extended Data Fig. 2).

**Fig. 1.**
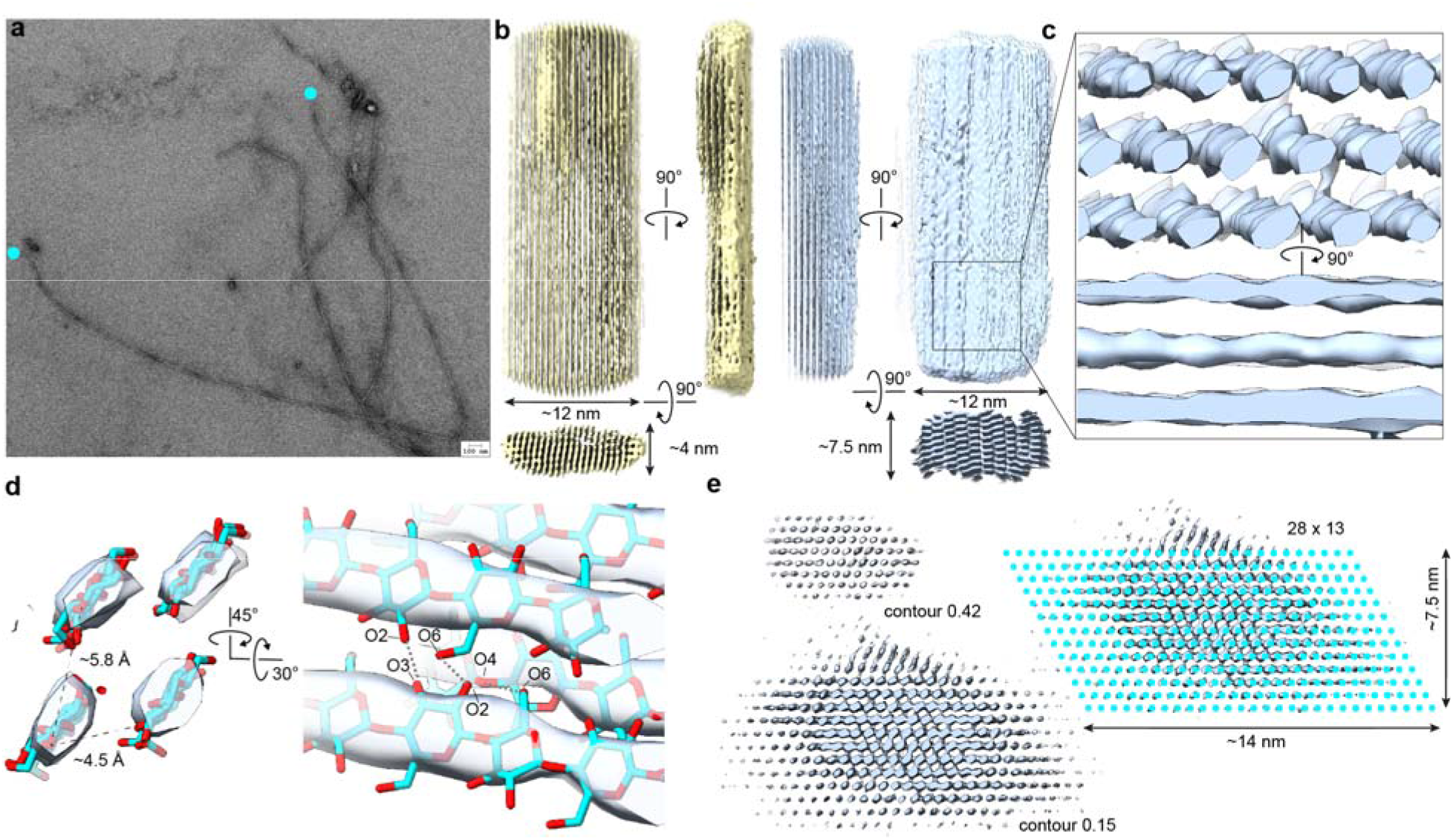
*Ciona intestinalis* cellulose fibril reconstruction. (**a**) Negative stain image of tunicate cellulose fibrils. The ends of one fibril are indicated by cyan circles. (**b**) Cryo-EM maps of a thin (yellow) and thick (blue) fiber reconstructions. (**c**) Locally refined map of the thick fibril focusing on the area indicated by a black rectangle in panel B, contoured at 0.47σ. (**d**) Arrangement of four cellulose strands after docking into the density shown in panel C. Proposed intermolecular H-bonds are indicated as gray dashed lines. (**e**) The same map shown in panel C but at different contour levels. Right panel: Estimated number of cellulose strands indicated by cyan circles.

Ab initio three-dimensional volume reconstruction revealed two fiber structures overall, referred to as thin and thick fibrils, with dimensions of approximately 4×12 and 7.5×12 nm, respectively (Fig. 1b). The thick fibril is consistent with previously reported dimensions of tunicate cellulose fibers obtained from TEM images of thin tunicate sections ^9,22,23^. The map quality of the thick fibril was improved by focused refinement of a smaller segment, resulting in a map of approximately 4.8 Å resolution that was used to generate an atomistic model (Fig. 1c and Extended Data Fig. 2). The map resolves densities of individual strands with oscillating thicker and thinner regions that match the glucosyl repeat units of cellulose. The coin-shaped glucosyl densities also define the sugar orientations relative to the fiber axis. Other features, however, such as the glucosyls’ C6 hydroxymethyl groups are not resolved, leaving the exact register of the glucan chains unresolved.

### Architecture of the tunicate fibril

A fiber model was built by placing cellohexaose molecules into the locally refined thick fiber density. Initially, four neighboring cellulose strands were placed, resulting in a rhomboidal segment with two glucan strands facing each other via their glucopyranose rings, separated by about 4.5 Å, and two strands aligned with their hydroxyl groups, separated by approximately 6 Å (Fig. 1d). This initial segment was expanded to obtain a fiber model containing 36 cellulose strands. The obtained model fits equally well into the thick and thin fiber densities.

At a low contouring threshold, the cryo-EM map reveals weaker densities of additional glucan chains surrounding the well-resolved core (Fig. 1e). This suggests a fiber containing approximately 28 chains laterally and 13 layers vertically (364 total), giving a dimension of 14 by 7.5 nm, consistent with earlier estimates ^23^. A similar analysis of the thin cellulose fiber suggests an arrangement of 6 × 22 glucan chains (Extended Data Fig. 3a).

Despite lacking insight into the register of the glucan chains, the fiber model was built such that glucans in neighboring layers are shifted along the fiber axis by about half a glucosyl unit, as also observed in crystalline cellulose ^19,20^. This arrangement places the C6 hydroxyl groups of one strand within H-bonding distance to the O4 oxygens of a stacking neighboring strand (Fig. 1d). Similar H-bonding distances exist in the other dimension. Here, intermolecular H-bonds are possible between the C2 and C3 hydroxyl groups of one chain and the C6 and C2 hydroxyl groups of another chain (Fig. 1d). These distances appear as connections in some map segments (Extended Data Fig. 3b).

In our fiber model, the glucopyranose rings of diagonally opposed single cellulose strands are exposed to the bulk solvent (Fig. 2a). This fiber face is referred to as (200) based on the cellulose I_β_ indexing, with the other faces referred to as (1-10) and (110) ^24^. Within the (200) layers, the glucan strands are separated by about 5.5 Å, and alternating juxtaposed layers are approximately 7.2 A apart from each other (Fig. 2b). Consecutive layers are shifted by ∼5 Å relative to each other along the (200) face, creating a running bond brick pattern of the fiber cross-section (Fig. 2a, b). The resulting fiber model exhibits slight electronegative and –positive surface properties (Fig. 2c).

**Fig. 2.**
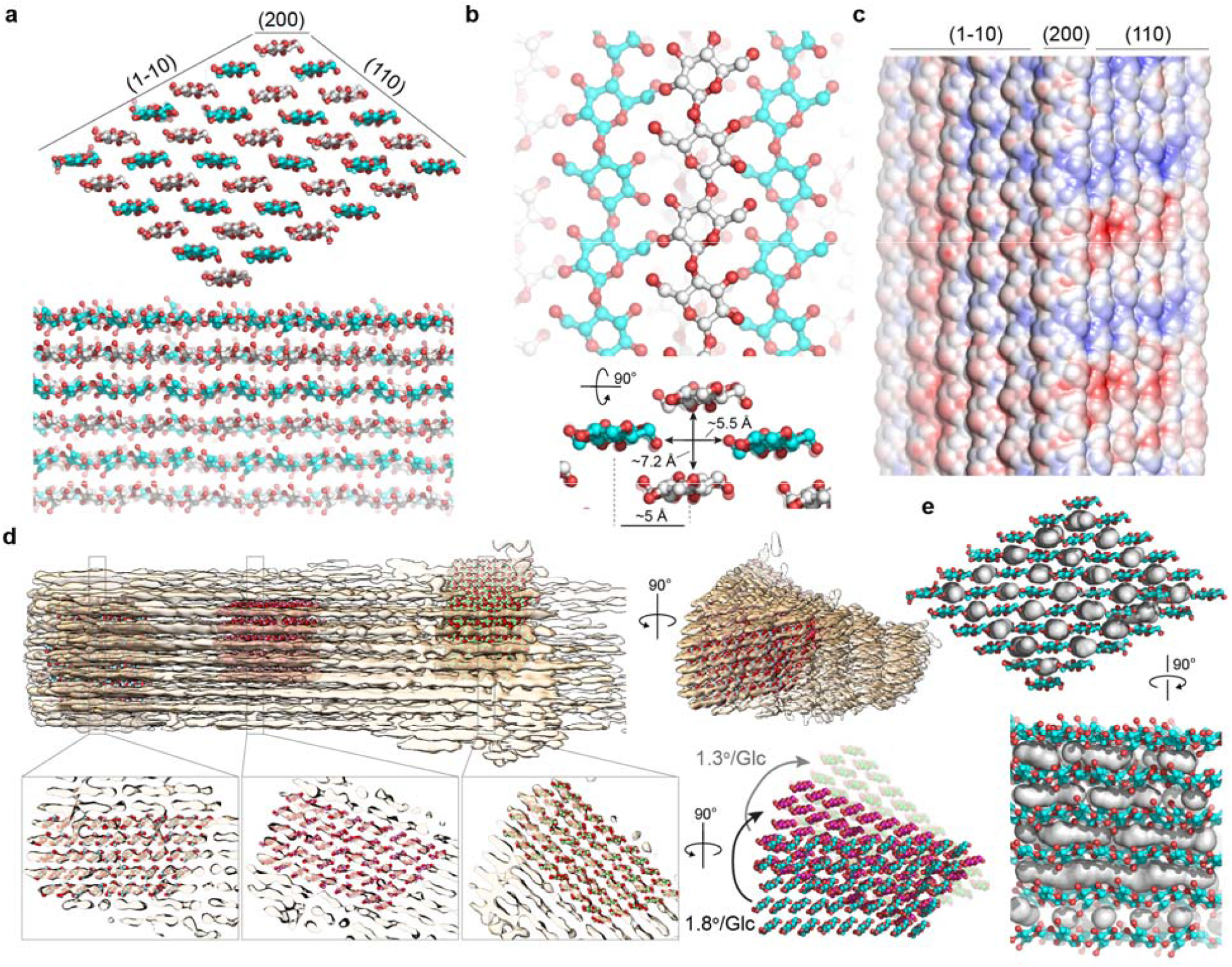
Architecture of the tunicate cellulose fibril. (**a**) Fiber model shown as a cross-section (top) and side view (bottom). Glucan stands belonging to different layers are colored in cyan or white for their carbon atoms. For orientation, fiber faces are annotated according to cellulose I_β_ convention. (**b**) Detailed view onto the (200) face of the fiber. (**c**) Electrostatic surface potential of the fiber calculated using the CHARMMGUI PBEQ solver ^54^. (**d**) Cryo-EM map of a twisted fiber with the model docked into it at the end and middle sections. (**e**) Solvent tunnels (gray surfaces) inside the fiber calculated using Pymol ^55^ with a 1.4 Å solvent probe.

### Twisted cellulose fibers

To reconstruct the volumes described above, fiber segments with a detectable twist were omitted. However, during refinement, a volume segment with a notable rotation around the fiber axis was obtained that resolved individual fiber layers (Fig. 2d). The map is of sufficient quality to dock the generated fiber model into it.

Comparing the models docked at one volume end and the middle section reveals a rotation of about 21.6 degrees over a distance of 62 Å. Similarly, the model docked at the opposing fiber end is rotated relative to the center position by 22.1 degrees, over an estimated distance of 88.4 Å. Considering a length of 5.2 Å per glucosyl unit suggests a fiber rotation of 1.8 and 1.3 degrees per glucosyl unit within the volume halves. More severe fiber twisting was frequently observed by visual inspection of cryo-EM images (Extended Data Fig. 1).

### The cellulose fibril contains internal water molecules

The separation of glucan chains within the (200) layer prevents direct H-bonds between neighboring glucan chains (Fig. 2a, b). This space is capped above and below by strands in neighboring layers, creating a solvent accessible volume at the interface of four glucan strands (solvent tunnels) (Fig. 2b, e). Therefore, we wondered whether the cellulose fiber is stabilized by internal structural water molecules.

To this end, we performed solid-state NMR spectroscopy of the never-dried fibers. Quantitative ^13^C NMR spectra show two sets of ^13^C chemical shifts that match the known I_β_ and I_α_ cellulose allomorphs ^25^, at an intensity ratio of 3:1 (Fig. 3a). Each allomorph resolves two C1 and C4 signals, likely corresponding to two inequivalent chains in each fiber. A small amount of disordered cellulose signals is also observed.

**Fig. 3.**
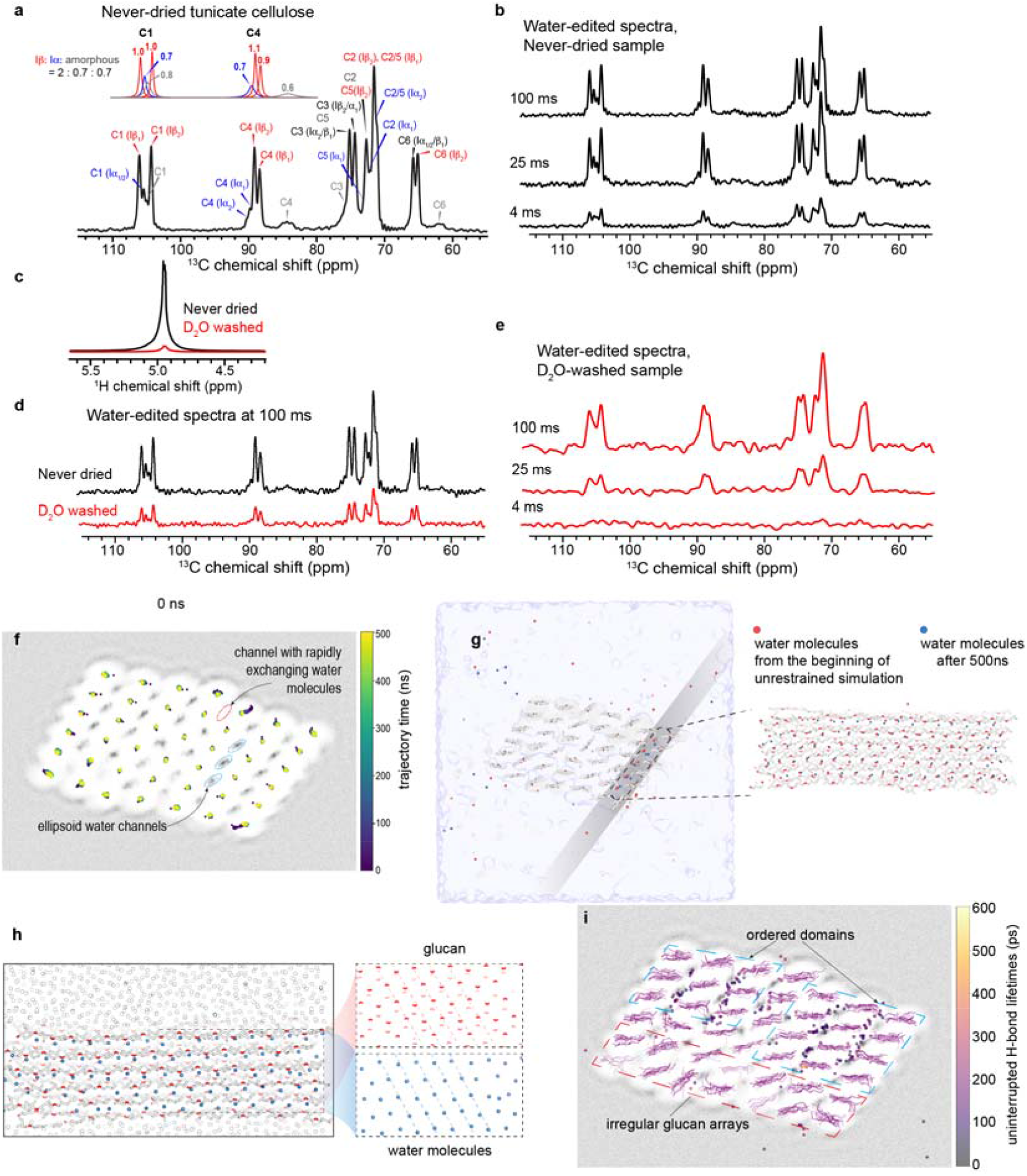
Hydration of the tunicate fiber. (**a**) Quantitative ^13^C NMR spectrum measured using multi-cross-polarization (multi-CP) experiments. Chemical shifts are assigned based on the literature ^45^. Mixed peaks are labeled black. (**b**) Water-edited ^13^C NMR spectra of never dried tunicate cellulose at different ^1^H spin diffusion mixing times. (**c**) ^1^H NMR spectra of the never-dried sample with H2O (black) and the D2O washed (red) sample. (**d**) Water-edited 13C spectra of D2O washed sample (red) compared to the original hydrated sample (black). (**e**) Water-edited ^13^C spectra of D_2_O-washed sample at varying ^1^H spin diffusion mixing times. (**f**) Transverse projection of the simulation box, with center of mass of ever glucan chain traced along the trajectory. Cumulative water positions are shown as a background greyscale heatmap (light to dark: increasing occupancy). (**g**) As in panel (F) but showing water occupancy as surface contour (blue) an cellulose strands as sticks. Water molecules present at the beginning and end of a 500 ns simulation are shown as spheres in orange and blue, respectively. Inset: Longitudinal cross section of a part of the fiber. (**h**) Longitudinal view after 500 ns simulation as in (g). Glucans are colored white with glycosidic linkages shown in red. Internal water molecules are shown as blue spheres. (**i**) Transverse projection view with water occupancy heatmap shown as in panel (f). Positions of glucosyl units are traced along the trajectory in greyscale (light-dark: 0-500ns). Average positions of waters mediating the longest uninterrupted H-bonds with glucosyl units are shown as a scatter plot, colored with the computed H-bond lifetime. Ordered and disordered fiber regions are indicated.

To probe the water accessibility of these cellulose fibrils, we measured water-edited ^13^C NMR spectra using variable ^1^H spin diffusion mixing times (Fig. 3b). The experiment selects the water ^1^H magnetization and transfers it to the glucosyl carbons in a time-dependent manner. Even at a short mixing time of 4 ms, the ^13^C spectrum of the native hydrated sample already shows 40% of the intensity of the 100 ms spectrum at which spin diffusion is complete. This suggests that the tunicate fibrils are accessible to highly dynamic water.

To determine whether some of the hydration water resides inside the fibril, we washed the never-dried sample with excess D_2_O, which depleted the total H_2_O pool to 8% of the original sample, as shown by the ^1^H NMR spectrum (Fig. 3c). Following this D_2_O wash, we remeasured the water-edited ^13^C spectra. Interestingly, the 100 ms water-edited spectrum retains ∼30% of the intensities of the unwashed sample at the same mixing time (Fig. 3d, e,) indicating that ∼30% of cellulose-bound water originates from the fibril interior, unable to exchange with external D_2_O.

### Structural water molecules stabilize four cellulose strands

To further corroborate the cellulose fiber hydration, we employed molecular dynamics (MD) simulations. To this end, a 42-chain fiber model with randomly placed water molecules inside the solvent tunnels was used. The simulation was set up with the cellulose fiber spanning the periodic boundaries of a cubic box of 96 Å, thereby generating an infinite fibril (Extended Data Fig. 4a). Following an initial equilibration for 100 ns with weak positional restrains (40 kJ mol^-1^ nm^-2^) on the glucan chains, five unrestrained 500 ns simulations were performed. The fiber remained stable during the trajectory, besides a slight flattening of strands relative to the (200) face, for unknown reasons (Extended Data Fig. 4a).

During the simulations, the solvent channels between the glucan chains remained solvated. Analyzing the positions of all water molecules in the simulation box reveals a finite volume for their movements inside the fibril (Fig. 3f). The water occupied volume resembles an elongated ellipsoid that runs diagonally between four glucan chains. While some internal water molecules near the fiber surface exchanged with bulk solvent, the majority of the internal water did not, or only slowly exchanged with bulk solvent, consistent with the ssNMR data (Fig. 3d, g).

The spacings between the (110) or (200) layers do not allow for an extensive H-bond network without water molecules bridging them. Such structured water should in turn also have a periodic arrangement similar to the periodicity exhibited by the glucans’ cellobiosyl repeat units. Accordingly, the water molecules in the same solvent tunnels adopt a periodic pattern with ∼5.2-5.6 Å spacing between them (Fig. 3h). Due to the register shifts between neighboring cellulose strands, the position of the structured water molecules also shifts by roughly 2.5 Å between layers. The thermal fluctuations predicted for these bridging water molecules are at the same levels as those of the nearby glucosyl moieties in the glucan chains, seemingly being held in place by the H-bonding hydroxyl groups (Extended Data Fig. 4b-e).

The fiber’s internal water molecules form H-bonds with the C2, C3, and C6 hydroxyl groups (Extended Data Fig. 4b). We speculate that water stabilization by the glucosyl units also stabilizes the corresponding cellulose strands. Indeed, the sugar-water H-bond pairs with the longest computed lifetimes do seem to regularize participating domains (Fig. 3I and Extended Data Fig. 4e).

Analyzing the mean glucan chain position over the simulation time revealed rearrangements of some surface exposed chains during the initial ∼200 ns, after which they remained stable (Fig. 3f, i). Further, some glucans exhibited a profound twist along their axis, suggesting that glucan chains within a hydrated fiber have some conformational freedom that does not destabilize the fiber architecture overall.

### Hydrated cellulose fibers share surface features with plant cellulose fibrils

We tested whether tunicate fibers interact with probes commonly used to visualize plant fibers. To this end, we probed fiber labeling with the cellulose-specific carbohydrate binding modules (CBM) 3a and 28, recognizing fibrillar and amorphous cellulose ^26-28^. Further, we also tested fibril binding by calcofluor, a common but less specific glucan probe ^29^, and analyzed fiber’s resistance to enzymatic degradation by glucosyl hydrolases. Plant cellulose fibrils are also known to interact with xyloglucan, a primary cell wall hemicellulose ^16^. This interaction was also probed using a xyloglucan specific CBM (CBM76) ^30^ for confocal and super-resolution MINFLUX fluorescence imaging ^31^.

As shown in (Fig. 4a-h), CBM3a conjugated to GFP or Alexa Fluor (AF) 647 readily interacts with the tunicate fibers, while AF660-conjugated CBM28 does not. Like CBM28, a hyaluronan-specific CBM (CBM70) ^32^ also does not interact with the cellulose fibers, while the small molecule calcofluor does, as expected. To test xyloglucan binding, the fibers were incubated with tamarind xyloglucan overnight, prior to labeling with a Flux 660-conjugated CBM76 and CBM3a-GFP. No CBM76 binding was observed in the absence of xyloglucan. In its presence, however, CBM76 efficiently detects the fibers. As a control, imaging xyloglucan alone with CBM76 or CBM3a in the absence of cellulose fibers only shows scattered background signals (Fig. 4g and Extended Data Fig. 5a).

**Fig. 4.**
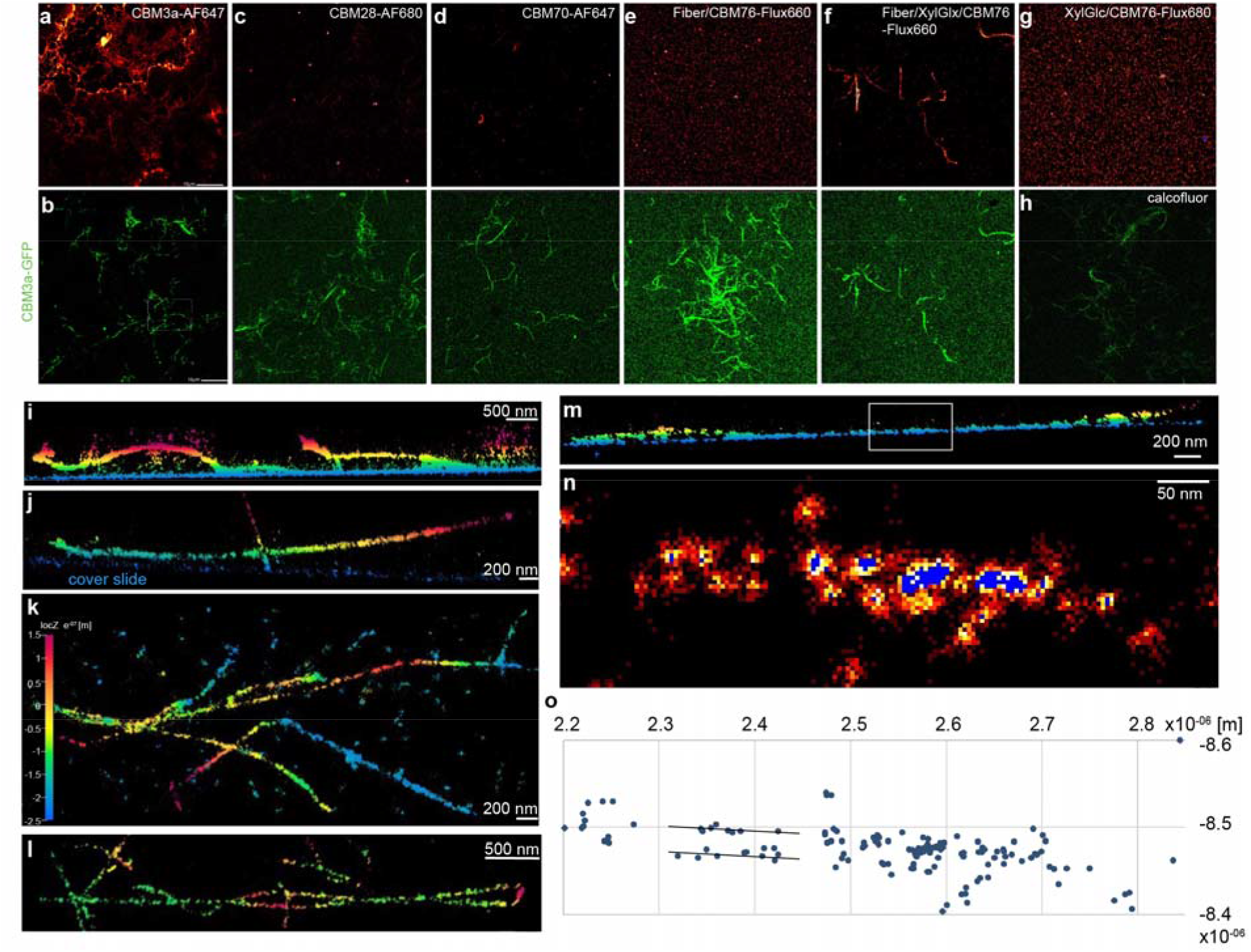
Tunicate and plant cellulose fibers have similar surface properties. (**a-g**) Confocal fluorescence images of tunicate fibers labeled with the indicated carbohydrate binding modules (CBM). In (c-f), the fibers were visualized with GFP-conjugated CBM3a as a control (bottom row). (g) No fibrillar structures are detected in the presence of CBM76 and xyloglucan only. (**h**) Calcofluor staining of the tunicate fibers. (**i-l**) Minflux nanoscopy of xyloglucan/CBM76-Flux660 labeled fibers. Background fluorophore binding to the glass support generates a thin blue line at xz or yz displays. Coloring indicates the distance in the z-direction away from the glass slide. (**m**) A linear fiber segment that is enlarged in panel (**n**) showing fluorophore localizations on presumably opposing sides of the fiber. (**o**) Averaged fluorophore localizations for the region in panel n.

MD simulations of the tunicate fiber in the presence of either CBM3a or xyloglucan oligosaccharides suggest that the probes bind to the same surface of the tunicate fiber (Extended Data Fig. 5b,c), consistent with previous estimates for plant cellulose fibers ^33,34^. To test the MD-predicted interactions, we employed MINFLUX nanoscopy to localize the probes on the tunicate fiber surface. In the presence of xyloglucan and Flux 660-labeled CBM76, fluorophores can be localized along the fiber, revealing straight, curved, and even kinked segments (Fig. 4i-l and Extended Data Fig. 6). Some fibers lie flat on the coverslip surface, others exhibit profound wave patterns and associations with or spiraling around other filaments. Thus, the twisting of the fibers observed on the vitrified cryo-EM grids is not an artefact, but rather an intrinsic fiber property. While being flexible, the fibers also exhibit stiffness, enabling curving away from the coverslip into the bulk buffer (Fig. 4i, j).

Focusing on straight fiber segments bound to xyloglucan and labeled with CBM76, the fluorophore localizations are aligned in two rows along the fiber axis in some regions, separated by approximately 28 nm (Fig. 4m-o). This distance is consistent with an ∼15 nm wide fiber associated with xyloglucan bound to a fluorescently labeled CBM approximately 4 nm tall. Analyzing CBM3a-bound fibers revealed a similar fluorophore distribution (Extended Data Fig. 6a-c), suggesting that both probes preferentially interact with likely opposing fiber surfaces. To test whether xyloglucan and CBM3a occupy the same fiber surfaces, a competition experiment was performed. Saturating the fibers with GFP-conjugated CBM3a substantially reduced or eliminated xyloglucan/CBM76 binding to the fibers (Extended Data Fig. 6d).

Lastly, plant cellulose fibers exhibit a remarkable resistance to enzymatic degradation, spurring the evolution of sophisticated degradation machineries ^35^. Accordingly, we tested fiber susceptibility to degradation by exo and endo β-1,4 glucanases ^36,37^. While negative stain EM imaging demonstrates enzyme binding to the fiber, only marginal roughening of the fibers was observed over seven days (Extended Data Fig. 7). However, fluorescent labeling of the released reaction products followed by polyacrylamide gel electrophoresis ^38^ identified released mono- or disaccharide units, suggesting that the fibers are not inert to degradation (Extended Data Fig. 7b).

## Discussion

Cellulose-producing organisms function in hydrated states, necessitating cellulose to form interaction networks in aqueous environments. Therefore, structural water molecules intersecting the cellulose fiber are likely a consequence of fibrillogenesis, similar to protein and nucleic acid folding in aqueous milieus ^39,40^.

Our MD simulations indicate that every glucosyl unit of the cellulose fiber mediates three intermolecular interactions on average, either to water molecules or to neighboring glucan chains. A seven-micron long fibril of 360 glucan chains would contain 1.3 × 10^4^ and 4.8 × 10^6^ glucosyl units per chain and fibril, respectively. This suggests that the fiber is stabilized by roughly 1.4 × 10^7^ intermolecular polar interactions, alongside van der Waals interactions between glucopyranose rings and entropic effects from solvent displacement during fibrillogenesis.

The cellulose I_α_ and I_β_ structures determined by diffraction methods on dried cellulose crystallites provided important insights into cellulose’s crystalline forms (Extended Data Fig. 8) ^19,20^. Contrasting the rigid crystallites, atomic force microscopy data of hydrated cell wall segments revealed an interweaving pattern of flexible fibers ^41,42^. Aligning a cellulose I_β_ model with our hydrated cellulose fiber structure suggests that solvent removal alone could account for the closer packing of the glucan chains in the crystalline material (Fig. 5a and Extended Data Fig. 8). Attempts to replicate the crystalline fiber architecture by drying tunicate fibers on cryo-EM grids produced fibers of similar morphology as the solvated sample, but the resulting map was of insufficient quality to confidently identify glucan strands or layers (Extended Data Fig. 9a-c).

**Fig. 5.**
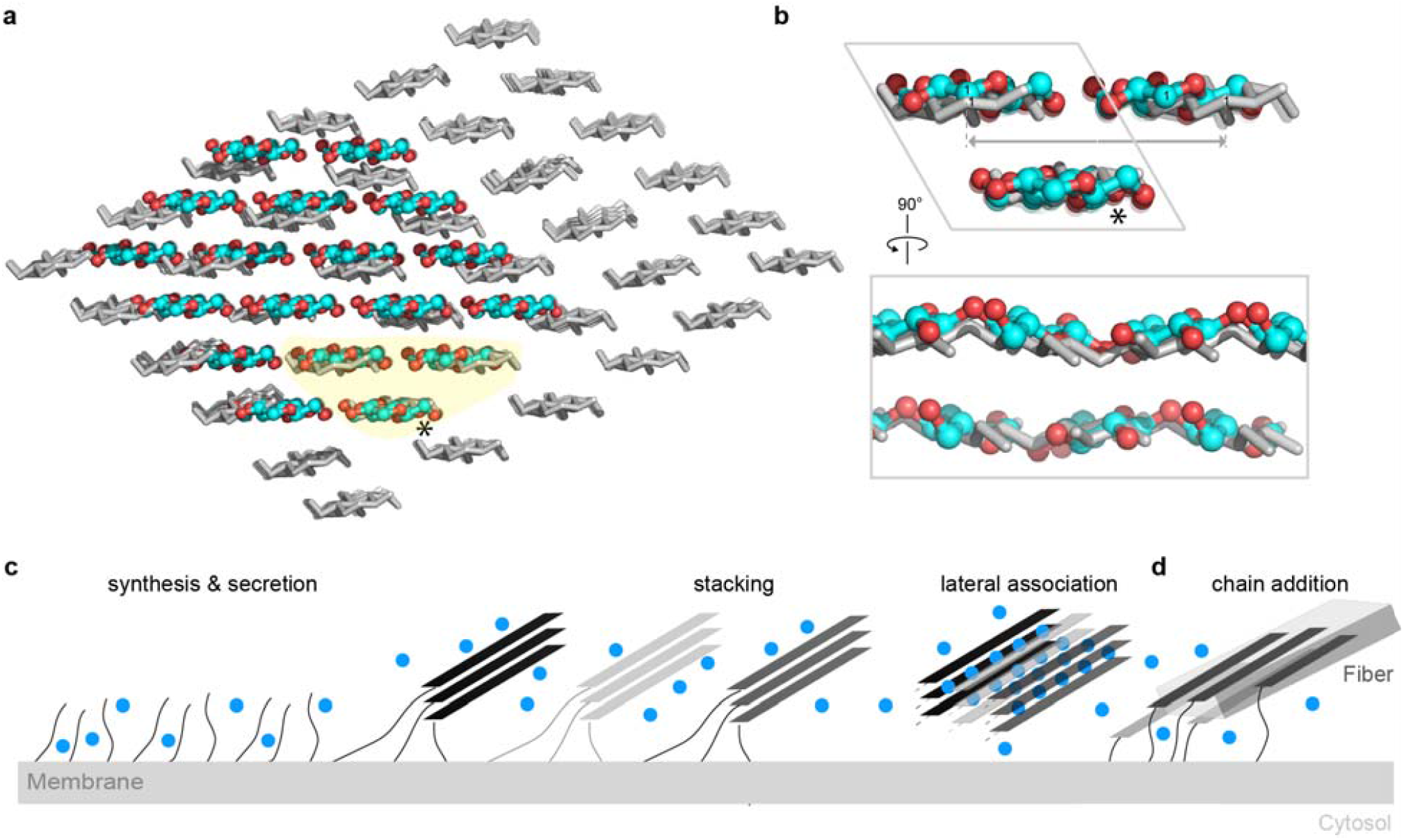
Comparison of hydrated and dehydrated cellulose fibers. (**a** and **b**) Superimposition of an 18-chain cellulose I_β_ model (colored cyan and red) with the hydrated fiber model (colored gray). The models are aligned based on the strand labeled with an asterisk and the shaded region is enlarge in (b). (**c**) Proposed model of fibrillogenesis in an aqueous environment. Individually secreted glucans likely stack via their glucopyranose rings. Stacks can than associate laterally, thereby trapping structural water molecules at the interface. (**d**) Large tunicate cellulose fibrils may arise from biosynthesis hotspots that add additional strands to a nascent fiber (indicated by a box). Water molecules are indicated as blue circles.

While high-resolution maps are not available for dehydrated cellulose, ^13^C ssNMR data provided insights into the structures of these dried fibers. Interestingly, dehydrated cellulose shows the same ^13^C chemical shifts as native hydrated cellulose, except with broader linewidths, indicating that the average conformation of dehydrated cellulose is similar to the hydrated cellulose, but with higher static disorder (Extended Data Fig. 9d-f). Rehydration restored the narrow linewidths. Moreover, water-edited NMR spectra showed the presence of internal water molecules after washing in D_2_O, similar to native cellulose, suggesting that drying generated a porous fiber that is readily occupied by water molecules upon rehydration (Extended Data Fig. 9f). It remains to be determined whether tunicates produce hydrated and unhydrated cellulose fibers or whether the observed structural differences between the crystalline and hydrated forms are due to different preparation methods. Similarly, fiber diffraction studies of chitosan revealed hydrated and unhydrated states, with a proposed distribution of water molecules inside the fiber similar to the tunicate cellulose fiber ^43,44^ (Extended Data Fig. 9g).

Our tunicate fiber ^13^C ssNMR spectra suggest the presence of cellulose I_α_ and I_β_ based on previously reported chemical shifts ^45^. Chemical shifts are sensitive to subtle differences in local torsion angles. Although the fiber architecture determined here does not reflect the crystalline structures, at the current resolution, our model is consistent with the stacking of glucan strands in the (200) direction (Fig. 5b). Detailed calculations of different fiber models may be required to identify the exact local conformations that account for the characteristic chemical shifts.

Plant cellulose microfibrils are likely generated by pseudo-sixfold symmetric assemblies of trimeric cellulose synthase complexes ^46^, producing six 3-stranded protofibrils that may coalesce into an 18-chain fibril ^15,47^. MD simulations set up to test this model suggest that (a) three hydrated glucan strands indeed rapidly and stably stack via their glycopyranose rings to form a protofibril, (b) six protofibrils quickly stack into sheets that can ‘fold over’ to form fiber-like lateral arrangements, and (c) lateral protofibril arrangements trap water molecules at the interface of the cellulose strands, similar to the water distribution in the tunicate fiber (Fig. 5b and Extended Data Fig. 10). Lastly, an 18-chain fiber model of the tunicate fibril is stable over a 500 ns simulation trajectory, suggesting that smaller cellulose microfibrils may also be stabilized by structural water (Extended Data Fig. 10a), as previously proposed ^48^. The dimensions of the hydrated 18-chain model are like those estimated for plant cellulose microfibrils ^49^. At interfaces, water molecules can adopt preferred positions with energy barriers hindering their exchange. These hydration forces may influence the macroscopic properties of the fiber, as previously formalized in the hygroelastic theory ^50^.

On a molecular level, little is known about how tunicate cellulose synthases are clustered and positioned to enable fiber formation. Seminal freeze fracture transmission EM analysis of *Metandrocarpa uedai* suggest multiple cellulose synthase terminal complexes (of unknown stoichiometry) surrounding a nascent cellulose fiber ^51^. Structurally, the enzyme is related to bacterial cellulose synthase but harbors a large N-terminal cytosolic extension and an extracellular glycoside hydrolase (GH6) domain (Extended Data Fig. 10d) ^10^. Recent studies on *Gluconacetobacter xylinus* cellulose synthase postulated a cytosolic filamentous ‘belt’ that may position cellulose synthases along the cell axis to promote fibrillogenesis ^52,53^. In tunicates, cortical cytoskeletal elements may perform a similar role. Further detailed structural analyses will be required to reveal the organization of these cellulose synthase complexes.

## Acknowledgments

We thank Daniel Cosgrove for providing the CBM76 expression construct and appreciate discussions with Ed Egelman and Ravi Sonani on fiber analysis and power spectrum interpretation. We are also grateful to Michael Purdy and David Cooper from MEMC for assistance in cryo-EM data collection and Jessica Matthias from Abberior for advice on MINFLUX data collection and processing.

## Funding

This work used the Titan Krios microscope in the Molecular Electron Microscopy Core which is supported by the University of Virginia School of Medicine, Research Resource Identifiers (RRID):SCR_019031. The Titan Krios (S10-RR025067) and K3/GIF (U24-GM116790) were purchased in part or in full with the designated NIH grants. M.H. and Y. P. were partially supported by NIH grant GM159321, awarded to M.H.. R.H. and A.A. were supported by NIH grant R35GM144130 awarded to J.Z.. C.L., L.W. and J.Z were supported by the Howard Medical Institute of which J.Z. is an investigator. K.N. thanks Nori Satoh for his long-standing support and JSPS KAKENHI (19KK0388) for funding.

This article is subject to HHMI’s Immediate Access to Research policy, which requires that this article be made publicly available as initial and revised preprints deposited on a designated preprint server under a CC BY 4.0 license.

## Author contributions

R.H. and J.Z. conceptualized the project. K.N. purified the tunicate fibers. R.H. performed all cryo-EM and fluorescence imaging experiments. J.Z assisted in cryo-EM data processing. A.A. assisted in performing MD simulations and analyzed the data. C.L. purified the CBM probes and assisted in fluorescent microscopy analysis, and L.W. assisted with PACE analysis. Y. P. measured the solid-state NMR data and M.H. and Y.P. analyzed the data. M.H. and Y.P. drafted the ssNMR part of the manuscript. J.Z. drafted the initial manuscript and all authors edited it.

## Data, code, and materials availability

Coordinates and the corresponding EM map have been deposited at the Protein Data Bank under accession code XXX and YYY.

