## Supplementary Information for "Electron microscopy reveals water networks inside hydrated cellulose fibers"

Material and Methods
Extended Data Figure 1-10

### Materials and Methods

#### Ciona cellulose fiber purification

Adult specimens of *Ciona intestinalis* type A (*Ciona robusta*) were provided by the Yutaka Satou laboratory (Kyoto University) via the National BioResource Project, Japan [56]. After anesthetization with 0.02% (w/v) Tricaine (T0941, TCI) in seawater, the tunics were surgically collected and minced. Tunic pieces (20 g) were treated twice with 0.5 L of 2% (w/v) KOH (168-21815, FUJIFILM Wako) under gentle stirring at 20°C for 6 h each, followed by washing with distilled water until neutral. To prevent beta-elimination, mild oxidation was performed at pH 4.8 [21]. The washed pieces (2 g) were finely chopped and suspended in 0.1 M acetate buffer (100 mL, pH 4.8) containing 0.1 mmol 4-Acetamido-TEMPO (350-45631, FUJIFILM Wako) and 10 mmol sodium chlorite (014265, Alfa Aesar) in an airtight flask. The reaction was initiated by adding 0.5 mL of 2M sodium hypochlorite (195-17212, FUJIFILM Wako). The flask was immediately sealed and stirred gently at 40 °C for 40 h. After cooling to room temperature, the suspension was thoroughly washed with milli-Q water via glass filter filtration (GC-50, Advantec). The oxidized cellulose was suspended in water at 0.05% (w/w) and mechanically dispersed using a homogenizer (Physcotron MS-56, Microtec Co.,Ltd.) at 7,500 rpm for 8 min. The dispersed nanofibers were collected by centrifugation (Avanti HP-20XP, Beckman Coulter) at 12,000 rpm for 20 min and stored at 4 °C.

#### NMR sample preparation

We prepared five cellulose samples for solid-state NMR measurements: 1) a never-dried sample; 2) a never-dried sample that was washed with $D_2O$; 3) an ethanol-dried sample; 4) an ethanol dried and then $H_2O$-rehydrated sample; and 5) a rehydrated sample that was washed with $D_2O$. These samples were used to investigate the water accessibility of tunicate cellulose and to distinguish internal from external water.

To prepare sample 1, we centrifuged 2.4 mL of tunicate cellulose suspension for 1 h at 55,000 rpm and 4°C to obtain an ~80 mg wet pellet, which was then spun into a plastic insert for the 4 mm MAS rotor. To prepare the $D_2O$-washed sample 2, we unpacked sample 1 from the rotor, suspended the pellet in 0.6 mL of 99% $D_2O$, incubated it for 15 min at 4°C, then centrifuged it at 55,000 rpm for 40 min. This resuspension-incubation-ultracentrifugation cycle was repeated twice, then the hydrated pellet was packed into the rotor for NMR experiments.

To prepare the ethanol-dried sample 3, we centrifuged 2.4 ml of native cellulose suspension to obtain ~104 mg of a wet pellet, then added 0.3 mL of 95% ethanol, incubated the mixture for 2 min, then dried the pellet with air at room temperature for 1 h. We repeated this ethanol soaking and drying cycle six times, giving a ~4 mg dry pellet. This dry mass thus indicates a hydration level of 96% for the original suspension. The dry material was cut with a razor blade into sections of a few mm each and packed into a 4 mm MAS rotor.

After completing the NMR experiments on sample 3, we resuspended the dry material in 1.5 mL deionized water and incubated it for 17 hours at 4°C. The sample was then centrifuged at 55,000 rpm for 24 h to obtain a 65 mg wet pellet, which resembled the never-dried pellet. This sample 4 was packed into a 4 mm rotor. After the sample 4 experiments were completed, we washed the sample with $D_2O$ following the same procedure as used for the never-dried sample to obtain sample 5.

#### Solid-state NMR experiments

All solid-state NMR experiments were carried out on a 400 MHz (9.4 T) Bruker AVANCE III spectrometer using a 4 mm $^{1}H/^{13}C$ MAS probe. Samples were spun at 7000 Hz at a sample temperature of 280 K. $^{13}C$ chemical shifts were externally referenced to the adamantane $CH_2$ peak at 38.48 ppm on the tetramethylsilane scale. Typical radiofrequency (rf) field strengths were 50-71 kHz for $^{1}H$ and 50 kHz for $^{13}C$. $^{1}H$-$^{13}C$ cross-polarization (CP) contact time was 1 ms and a linear ramp of 70-100% was applied on the $^{13}C$ channel. Recycle delay was 2 s.

To obtain quantitative intensities that reflect the relative concentrations of different carbons in this natural abundance material, we conducted multiple cross-polarization (multi-CP) NMR experiments [57]. The $^{1}H$-$^{13}C$ CP contact times and $^{1}H$ spin-lattice relaxation periods $t_z$ were optimized using a $^{13}C$, $^{15}N$-labeled glutamine sample. The optimized conditions include a $t_z$ of 0.9 s, a CP contact time of 550 µs, a 90-100% $^{13}C$ ramp, and six cycles of this module. The recycle delay was 2 s. $^{13}C$ DP spectrum with a 35 s recycle delay verified the multi-CP spectra of glutamine to be quantitative.

Water-edited $^{13}C$ CP experiments [58] were conducted using a 1.14 ms water $^{1}H$ $T_2$ filter containing a $^{1}H$ Gaussian 180° pulse of 1 ms and $^{1}H$ spin diffusion mixing times of 4, 25, and 100 ms. Because the samples are natural abundance in $^{13}C$, these water-edited spectra were signal-averaged with 12,000-96,000 scans to obtain sufficient spectral sensitivity.

**EM grid preparation and data collection**

The purified *Ciona intestinalis* cellulose fibers were serially diluted and 4 µL of each dilution was applied to a glow discharged Formvar/Carbon grid (Electron Microscopy Sciences) for 30s, followed by two washes with 4 µL double distilled (MilliQ) $H_2O$. The grid was negatively stained with 4 µL 0.75% Uranyl Formate (UF) in $H_2O$ for 30s. Excess UF was removed by blotting with filter paper, and the grid was air dried. Images were taken on a Tecnai F20 at the Molecular Electron Microscopy Core (MEMC) facility at the University of Virginia.

For cryo grid preparation, about 3 µL of the fibers at different concentrations were applied to C-flat 300 mesh 1.2/1.3 copper grids (Electron Microscopy Sciences), glow-discharged in the presence of amylamine at 25 mA for 45 s, back blotted with a Leica GP2 for 10 s at 4°C, 100% humidity, and then frozen in liquid ethane.

Cryo-EM data were collected at MEMC on a Titan Krios microscope operated at 300 keV and equipped with a Gatan K3 direct electron detector positioned post a Gatan Quantum energy filter. Total 8,147 movies were collected in counting mode at a magnification of 105,000, calibrated pixel size of 0.652 Å, and defocus range from -2.0 to -1.0 µm with step size of 0.2 µm. The total dose was 50 $e^-/Å^2$. Movies with 40 frames were collected at 5.17 s/movie rate.

**Cryo-EM data processing**

Data processing was performed in CryoSPARC v4.7.1 [59]. Movies were imported and patch motion corrected, followed by patch CTF estimation. The obtained micrographs were curated based on relative ice thickness, estimated resolution, defocus range, and full frame motion distance. Particles were picked in two rounds. First, template-free filament tracing using a 180 Å filament diameter, a separation distances between segments of 0.2, and a minimum filament length of twice the diameter. Further, hysteresis low and high thresholds of 89 and 96% were applied, respectively. The selected particles were extracted and processed as described below to generate templates for template-based particle picking.

Template based particle picking was performed using the filament tracer with 28 2D class averages as templates, a filament diameter of 180 Å, a separation distance between segments of 0.3, and a minimum filament length of 2 diameters to consider. The low and high hysteresis thresholds were 89 and 96%, respectively. The inspected picks were extracted with a 600 pixels box size and 4-fold Fourier cropped to a box size of 150 pixels, followed by multiple rounds of 2D classification as shown in (fig. S2). During this process, care was taken to omit 2D class averages of fiber segments with a detectable twist. The cleaned particle stack was used to generate five ab initio classes, followed by heterogenous refinement of all classes. This identified the thick and thin fiber classes, which were processed separately thereafter.

The thin fiber class was further refined by heterogeneous refinement using all volumes generated during the initial ab initio reconstruction, two rounds of 2D classification, non-uniform and local refinement with a solvent mask that covered the entire fiber segment, as well as 3D classification, also using a solvent mask. Finally, the particles were reextracted using a 300-box size (2x Fourier cropped), followed by non-uniform and local refinement using a solvent mask. Focused refinements using smaller masks at various locations of the fiber segment did not improve the map quality. The final map was generated using 80,081 particles.

The thick fiber was obtained similar to the thin fiber but required more sorting steps in two and three dimensions (fig. S2). The map quality improved significantly after applying a focused mask covering about 1/3 of the fiber segment. This mask was further refined and reduced in volume to generate the final map from 47,146 particles at an estimated resolution of about 5 Å. We noticed that resolution estimates fluctuated between 4.5 and 5.5 Å upon subtle changes to the focused mask, without apparent quality differences of the obtained maps. All maps were automatically B-factor sharpened in CryoSPARC.

**Model building**

Initially we attempted to dock an 18-stranded fiber model (containing cellohexaose strands) of cellulose $I_b$ into the cryo-EM map, without success. Thus, the glucan strands of the cellulose $I_\beta$ model were translated in the x/y dimensions perpendicular to the fiber axis as rigid bodies. Of the obtained model, a segment containing four glucan strands was used as a repeat unit to build larger cellulose fiber models by rigid body docking.

The individual glucan strands of the obtained model were real space refined in Coot [60] using a map refinement weight between 5 and 10. Following manual model building in Coot, the fiber model was real-space refined in phenix:refine [61,62] using a refinement weight of 0.5.
For molecular dynamics simulations, the fiber model was expanded to 18 glucosyl units per glucan strand. In addition, water molecules were placed randomly inside the solvent tunnels in Coot. Water placement was based on visual inspection of the solvent channels and potential hydrogen bonding with carbohydrate hydroxyl groups. The final model used for input file generation in CHARMM-Gui lacked the C1 hydroxyl group at the glucans' reducing end to generate a continuous fiber across periodic boundaries.

**MD simulations**

MD simulations were performed using a 42-chain cellulose fiber with 18 glucosyl units per strand and randomly placed water molecules in the fiber's solvent channel. All input files for Gromacs were generated using the Glycan Reader & Modeler feature of the CHARMM-GUI using the CHARMM36m carbohydrate force field [63,64]. Specifically, the glucan strands were defined as cyclic glycans with a b-(1,4) linkage between the first and last glucosyl unit. The fiber was placed in a cubic solvent box with 150mM KCl with a dimension of 96 Å. The box size was optimized based in test simulations using box sizes from 93 to 98 Å in 1 Å steps.

MD simulations were performed using the GROMACS 2025 package [65]. Energy minimization was done with steepest descent integrator till the emtol stabilized to 1000 kJ/mol.

One round of energy minimization with V-rescale thermostat was done for 1 fs timestep for 125 ps at 310 K. To optimize fiber stability, mildly restrained initial simulations were run for 100 ns introducing a semi-isotropically coupled C-rescale barostat with a timestep of 2 fs. During these runs the carbohydrate components of the system were restrained with 40 kJ $mol^{-1}$ $nm^{-2}$ while solvent molecules were unrestrained. The initial extended equilibration was followed by at least 500 ns of unrestraint simulation. The simulations were run with 5 independent random seeds. Similar simulations were also performed for a 3 chain and an 18-chain system.

Simulations including CBM3a or xyloglucan were also performed as described above. For CBM3a, the protein was place into the simulations box in three different starting poses, with residues predicted to interact with cellulose either facing the fiber, pointing away from it, or arranged perpendicular to the fiber axis. For the xyloglucan simulations, an oligosaccharide consisting of seven β-1,4 linked glucosyl units with a central unbranched glucose unit was generated using the glycam server (glycam.org) [66]. The first and the last three glucosyl units contained an α-xylosyl residues attached to their C6 hydroxyl groups (i.e. XXXGXXX).

For analyzing glucan chain motions in context of the entire fibril, the trajectories were first exported without hydrogens through gmx trjconv with -pbc mol flag, using the glucan chains for alignment. Since it was an infinite fibril, the pdb trajectory still showed translation of entire chains along the axis of the fiber out of the periodic boundary in some cases, and hence these jumps were aligned with the first frame using pymol. This input was used for calculating intrafibril motion of chains, and to calculate RMSF of water molecules. This may have given artefactual values for bulk waters due to rotational transformations on glucan chains during alignment but provided near identical RMSF values for waters trapped inside the fiber before and after alignment. For H-bond analysis and its effect on the participating residues, the trajectory after the aforementioned trjconv step was used for analysis, and the topology information was imported from the respective tpr files through the MD Analysis library package. H-bond lifetime calculations were done through the hydrogenbondanalysis module in the same library [67,68], with no intermittent H-bond breakage tolerance and 100 ps timesteps, giving 5000 datapoints for each interacting pair. The coordinates and atom info (i.e., atom, residue and chain indices, and atomtype) were extracted for regression analyses, while Hydrogen Bond Lifetimes (hbl) calculations were done using default parameters (donor-acceptor distance cutoff at 3Å, and donor-hydrogen-acceptor angle be greater than 150°) between the sugar hydroxyls and water molecules. The weakest interacting H-bonds did not display a first-order decay for hbl, and hence exponential decay curve fit did not work for the majority of weak interactions. Area under the curve was hence used as a fallback option. We found that at least for the interactions whose hbl could be fit with an exponential decay function, the corresponding H-bond lifetime constants closely matched with those calculated via the latter method. To correlate the propensity of H-bond interacting atom/sugar with the lifetime of the interaction, a very coarse correlation was performed, where the time averaged value for the root mean squared fluctuation (rmsf) of the atom or atom group, or the bond associated angles/torsion angles were calculated and correlated with the atom-index-associated hbl, if they were present in the top 5000 longest H-bond lifetime constants observed in the trajectory. To make sure that inherent physical restraints (such as O6-C6-C5-C4 dihedral angle that may have a split population away from the mean angle) seen in some observables do not affect our ability to assess them via just rmsf, the rate of such fluctuations were also calculated through autocorrelation function (acf), where the last significantly positive autocorrelation above 95% confidence interval along time axis was used as a measure for the pace of such fluctuations. To summarize, for every sugar atom that interacted with a water molecule through H-bonding, we checked the magnitude of fluctuation in the bonded neighborhood through rmsf, and its rate through acf. Where H-bond analysis was not performed, all analyses were performed using in-house scripts.

**Molecular visualization**
Fiber models were visualized and images were prepared in PyMOL, and ChimeraX [55,69]. The electrostatic properties of the cellulose fiber were calculated using the CHARMM-GUI PBEQ solver [54,70]. The fibrils in the MD simulations were arranged along the X-axis of the box and hence no further transformations were performed to generate YZ projection images. In cases where sliced views from particular views were needed, the rmsf/acf values were exported to the b-factor column of the first PDB frame of the trajectory and visualized through PyMOL [55].

**Synthesis and cloning of CBMs**

For the construction of CBM plasmids, the *E. coli* expression vectors pET28a-His$_6$-CBM3a-twinstrep and pET28a-His$_6$-twinstrep-CBM28 were synthesized (Gene Universal, Newark, USA). The CBM sequences used in this study correspond to the DNA fragments encoding CBM3a (amino acid residues 365-523 of the full-length CipA cellulosomal scaffoldin from *Clostridium thermocellum*; GenBank accession number CCV01464.1) and CBM28 (amino acid residues 561-752 of the full-length β-1,4-endoglucanase from *Ruminiclostridium josui*; GenBank accession number BAA12826.2). After synthesis, the SNAP domain [71] was fused to the C-terminus of each CBM, separated by a four amino acid long linker (GSGS), using the NEBuilder HiFi DNA Assembly Master Mix (NEB, MA, USA). In addition, CBM3a was also C-terminally fused to Emerald GFP (EmGFP) in the pET30a vector, without an additional affinity tag.

The CBM76 construct was a gift from Dan Cosgrove (Pennsylvania State University). The construct was PCR amplified using the following forward and reverse primers: Forward: tattaccatgggtgagaagatatacgaggaaaagaag, reverse: gtgatgagccgcagactaacctcagcttccgcCACCATCACCACCACCACCATCACCACtaactcgagtata. The gene was cloned using NcoI and XhoI restriction site into the pET30a vector, in frame with an existing SNAP domain, thereby generating a SNAP-CBM76- His$_9$ construct.

The SNAP-CBM70 construct was also cloned into the pET30a expression vector, as described before [72].

**Production and purification of recombinant CBMs**

Following sequence verification, all CBM constructs were transformed into *E. coli* BL21 (DE3). Glycerol stocks of the transformed strains were prepared and stored at -80 °C. These glycerol stocks were subsequently used to inoculate 100 mL of LB medium containing 50 µg/mL kanamycin in 200 mL flasks. This pre-culture (10 mL) was then used to inoculate 1 L of LB medium containing 50 µg/mL kanamycin in 2.5 L shake flasks. The flasks were incubated at 37 °C and 200 rpm until the culture density reached an optical density of 0.6~0.8. Protein expression was induced with 0.5 mM IPTG at 20 °C for 16 hours. Cells were harvested by centrifugation at 4,000 rpm for 15 min, cells were stored at -80 $^o$C until further use. The cell pellet was thawed, resuspended in a buffer containing 10 % glycerol, 100 mM NaCl, and 20 mM Tris (pH 7.5), and incubated with 1 mg/mL lysozyme for 1 hour. After adding 1 mM PMSF, the cell suspension was lysed by three passes through a microfluidizer (18 kpsi). The cell lysate containing the target protein was centrifuged at 200,000 g for 30 min in a Ti45 rotor (Beckman), and the insoluble fraction was discarded. The supernatant was supplemented with 20 mM imidazole and incubated with 5 mL bed volume of Ni-NTA resin for 1 hour at 4°C with agitation. The resin was washed sequentially with 1) 1 M NaCl in PBS (pH 7.4) containing 40 mM imidazole, and 2) PBS containing 60 mM imidazole. The target protein was eluted after a 30 min incubation in PBS containing 320 mM imidazole and concentrated to 1 mL using a 10 kDa molecular weight cut-off filter (Amicon). The concentrated sample was dialyzed against PBS overnight, aliquoted, flash-frozen in liquid $N_2$, and stored at -80 °C. The CBM3a-EmGFP protein was expressed and purified as described above with the exception that the protein was bound to Ni-NTA resin based on promiscuous binding, washed as described, and eluted with 100 mM glycine-HCl (pH 2.0), immediately neutralized with 1 M Tris-HCl (pH 8.0), and subsequently dialyzed against PBS pH 7.4.

**CBM-SNAP labeling**

All labeling steps were performed in the dark at 4 °C. Aliquots of the SNAP-tagged CBMs were thawed and mixed with DMSO-solubilized SNAP-surface AF647 (S9136S from NEB), SNAP-Flux660 (Abberior) or AF660 C2 Maleimide (A20343 from ThermoFisher) at a 1:1 molar ratio in the presence of 0.1 mM TCEP, respectively. The mixture was incubated under agitation for 6-8 hours. Size exclusion chromatography was performed using a Superdex 200 10/300 GL column equilibrated with PBS to separate the CBM-SNAP-AF conjugates from the unreacted free dye. Peak fractions exhibiting strong absorbances at 280 nm and 647 or 660 nm were pooled, adjusted to 2 mg/mL, aliquoted, flash-frozen in liquid, and stored at -80 °C.

**Minflux sample preparation**

High Precision glass coverslips (24mm, 1.5H. Marienfeld) were glow discharged at 25 mA for 90 s and each coverslip was coated with 100 mL poly-lysine and gold beads (25 mL 0.1% poly-L-lysine, 5 mL spherical gold nano particles (NAN PAR) and 70 mL PBS). After a 20 min incubation at RT in a damp chamber, the coverslips were rinsed and kept in PBS for later use.

**Fluorescent labeling of the tunicate fibers**.

For labeling for confocal imaging with CBM3a-AF647, CBM28-AF660, CBM70-AF647 and CBM76-flux660 (no xyloglucan): 10 mL fiber suspension in PBS (sonicated for 2 minutes in a cold water bath) were incubated with the fluorophores overnight in the dark at a concentration ranging from 30 nM to 7 µM, optimized for each fluorophore (to compensate for the different conjugating efficiency). Then, the sample was diluted 5-fold in PBS, applied to a coated coverslip, and incubated for 1 h at RT in the dark. The coverslip was rinsed twice with PBS and mounted in 10 mL PBS and sealed with nail polisher.

For staining with Calcofluor White Stain (Sigma) for confocal imaging: 50 mL fiber suspension in PBS was applied to a coated coverslip, incubated in a wet chamber for 1 h at RT, rinsed twice with PBS, and mounted in 5 mL PBS and 5 mL Calcofluor White Stain mixture (Sigma-Aldrich I8909), then sealed with nail polisher.

For staining of xyloglucan with CBM76 for confocal and Minflux imaging: 50 mL of the fiber suspension in PBS were incubated with 5 mL 1.0 mg/ml xyloglucan in 0.5 mM Na acetate, pH 5.5 (Megazyme, xyloglucan was prepared by sonication and boiling at 100 ºC for 10 mins at 1 mg/ml in 50 mM Na acetate) for 3 to 5 hours at 4 ºC. The fibers were then diluted based on initial imaging results, and CBM76-Flux660 was added at 30 nM, followed by overnight incubation in the dark. 10 mL of the sample was diluted 5-times in PBS, applied to a coated coverslip, and incubated for 1 h at RT in the dark. The sample was then rinsed with PBS and mounted in 210 mL of mounting buffer, containing: 50 mM Tris/HCl pH 8, 10 mM NaCl, 10% d-glucose, 100 U/mL glucose oxidase (Sigma), 1,200 U/mL catalase (Sigma), and 12 mM cysteamine (MEA) (Sigma). After blotting, the coverslip was sealed with a 1:1 mixture of the base and catalyst of Elite Double, 22 Regular Silicone Duplicating Material (Zhermack) and incubated in the dark until the seal was dry (~20 mins).

For fiber labeling with CBM3a-conjugated AF647 for Minflux imaging: The fiber suspension was incubated with 700 nM CBM3a in PBS overnight in the dark. 10 mL of the sample was diluted 5-times in PBS to prepare a coverslip and mounted as described above. [73]

**Minflux and confocal data collection and processing**

Confocal and Minflux data were recorded on a commercial Abberior Instruments Minflux setup (Abberior Instruments GmbH), similar to the one reported by Schmidt et al. [73]. The system was equipped with a 100x/1.45 NA magnification oil immersion lens, a 640 nm continuous-wave laser for exciting CBM3a-AF647, CBM28-AF660, CBM70-AF647 and CBM76-Flux660, a 488 nm pulsed laser for exciting CBM3a-GFP, and a 405 nm continuous-wave laser for activation. Minflux imaging was performed with the 3D imaging protocol provided by Abberior Instruments with a lower background threshold (lowbgc 2000~5000) and a 640 nm excitation power of 67 µW. The single molecule emission was detected between 650-720 nm, and the 405 nm power gradually increased throughout the measurement in the nW range to ensure a constant single molecule detection rate. Confocal images were collected with laser power of 57 mW and 20 mW for 488 nm and 640 nm lasers, respectively, a pixel size of 50 nm, a dwell time of 10 µs, and a detection window of 500-550 nm and 650-720 nm. All images were recorded with the pinhole set to 0.83 AU. The sample position was actively stabilized on the back-scattered light of gold beads illuminated with 980 nm in widefield mode. The Abberior Instruments Imspector software (v. 16.3) with MINFLUX drivers was used to operate the system.

**Minflux data processing**

In Excel, we first excluded the possible dual fluorophores events by filtering out the events with less than 10 or more than 206 iterations with the same trace ID (tid). Then the average of each fluorophore's parameters (per tid) were calculated. By comparing the averaged $7^{th}$ and $9^{th}$ iteration's detector channel ratio (dcr), the fluorophores were kept only when the difference of its $7^{th}$ and $9^{th}$ iteration's dcrs was less than 0.05 (this excluded the possibility of two fluorophores with different colors were located very close to each other and being excited at almost the same time). Finally, only the fluorophores with the dcr in the range of 95% confidence level of each fluorophore were assigned to the respective CBMs.

**Fiber digestion with glucanases**

The tunicate fibers were digested in a total volume of 40 mL of 50 mM $NH_4OAc$ pH 5.0, by adding 2.4 U of *Aspergillus niger* endo-β-1,4-glucanase (Megazyme) or $6x10^{-3}$ U *Trichoderma reesei* cellobiohydrolase-1 (Megazyme), and incubation at 37 °C for 3-4 days before refreshing the enzymes and incubation for a total of 7 days.

For PACE analysis [38], the samples were vacuum dried using a SpeedVac centrifugal concentrator (Savant). For labeling, the samples were resuspended in 10 μL ANTS master buffer, containing equal volumes of 15% acetic acid, DMSO, 0.2 M 2-picoline borane (2-PB), and 0.2 M ANTS (8-aminonaphthalene-1,3,6-trisulfonic acid) in 15% acetic acid, followed by incubation at 37 °C overnight in the dark. The samples were dried and resuspended in 10 μL of 6 M urea. Of this, 2.5 μL were loaded onto a 20% acrylamide PACE gel (240x180x0.75 mm, in 0.1 M Tris-borate pH 8.2), run first at 200 V for 30 mins, followed by 2 h at 1000 V in 0.1 M Tris-borate buffer using a Hoefer SE660 electrophoresis system. The gels were scanned using a Syngene G:box imaging station with a 365 nm transilluminator and 515-600 nm filter.

### Extended Data Figures

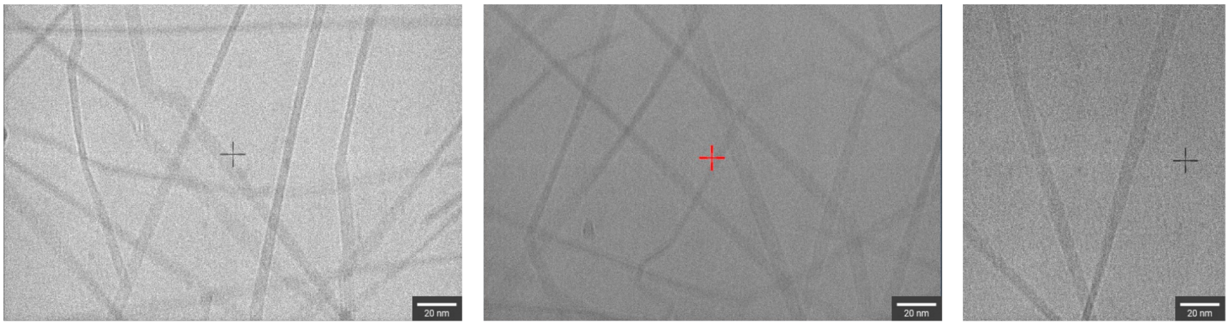


**Extended Data Fig. 1| Images of vitrified cellulose fibrils with profound twists along the fiber axes.**

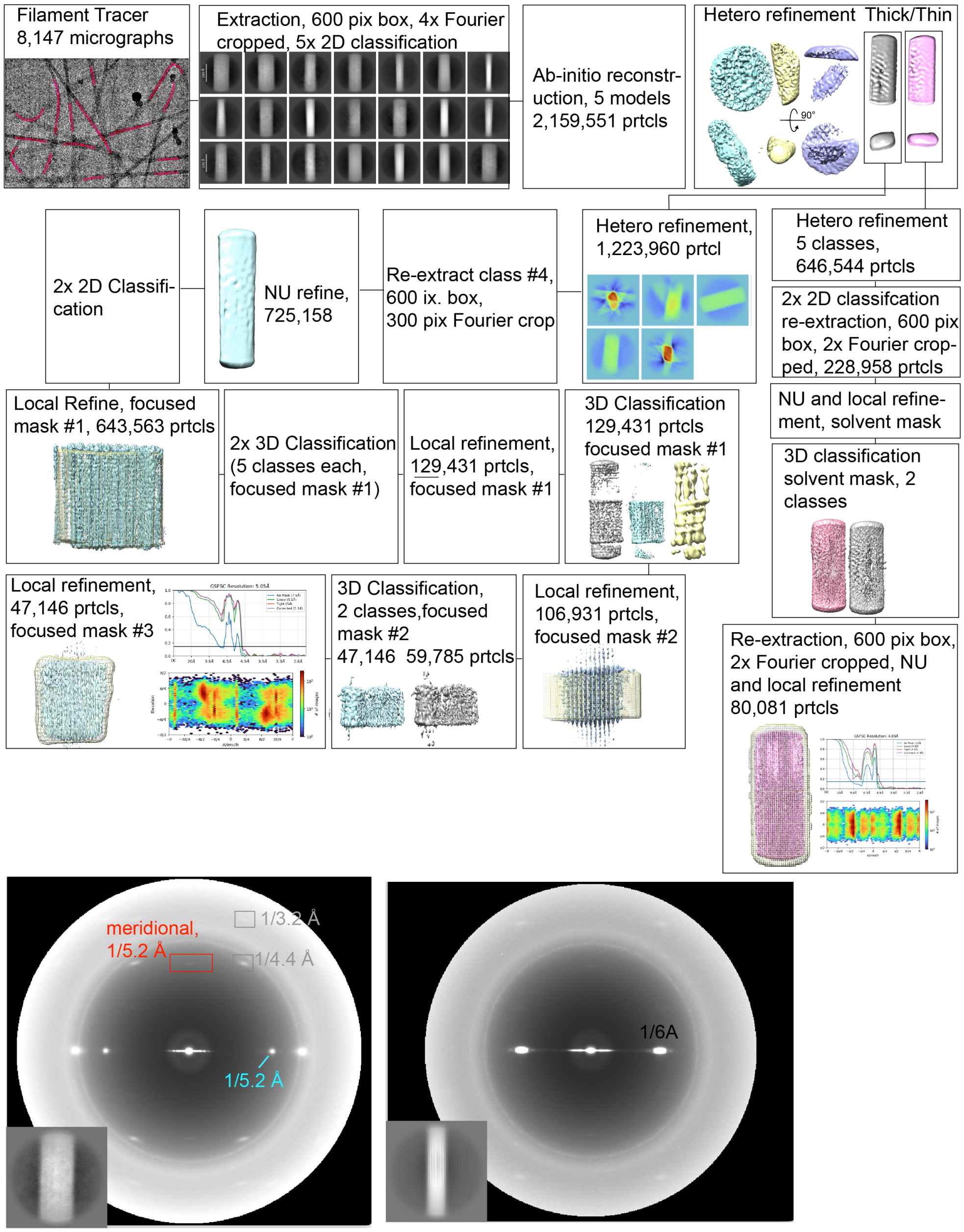


**Extended Data Fig. 2| Cryo-EM data processing workflow and power spectra of the indicated 2D class averages.**

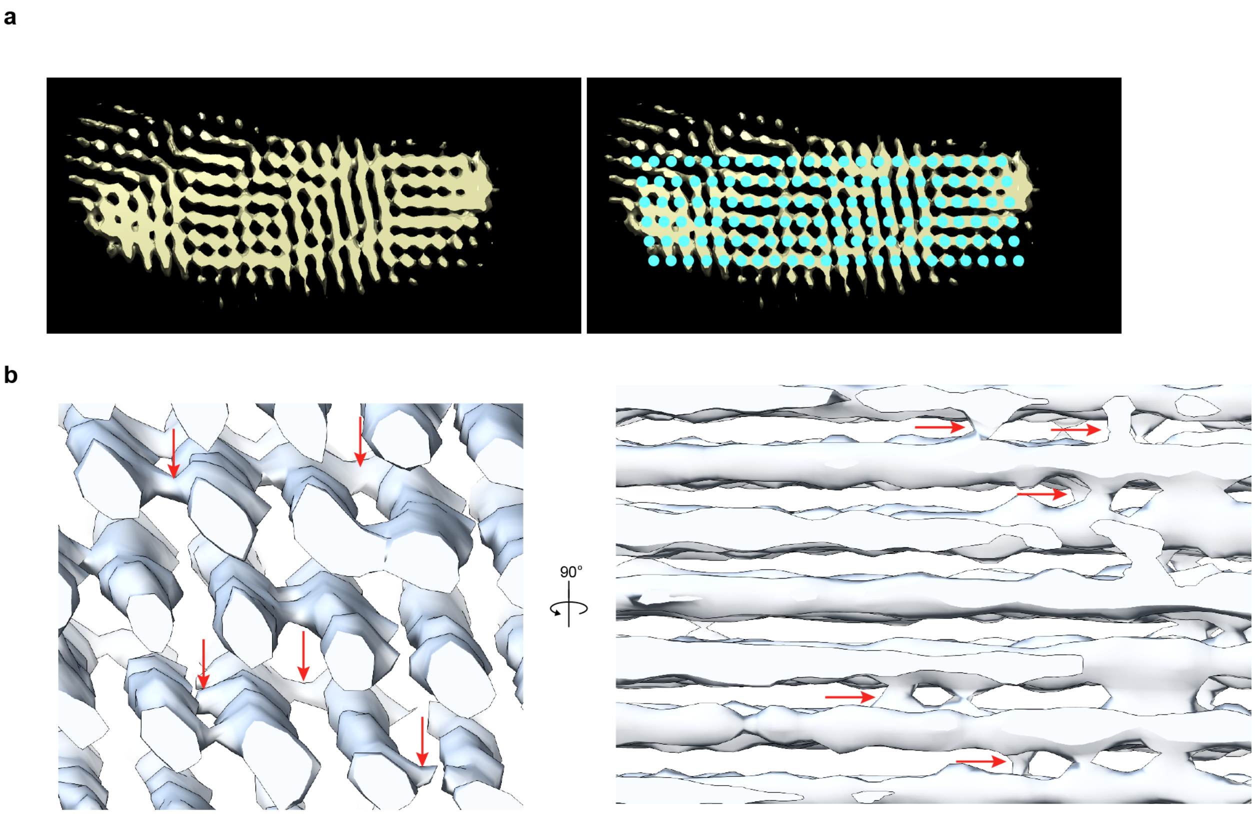


**Extended Data Fig. 3| Additional views of cryo-EM maps.** (**a**) Cryo-EM map of the thin fibril contoured at 0.12 σ. Cyan circles indicate suggested positions of glucan strands. The map also reveals fiber misalignment resulting in significant densities outside the core region at this contouring threshold. (**b**) Locally refined map of the thick fibril showing connections between neighboring glucan strands, indicated by red arrows.

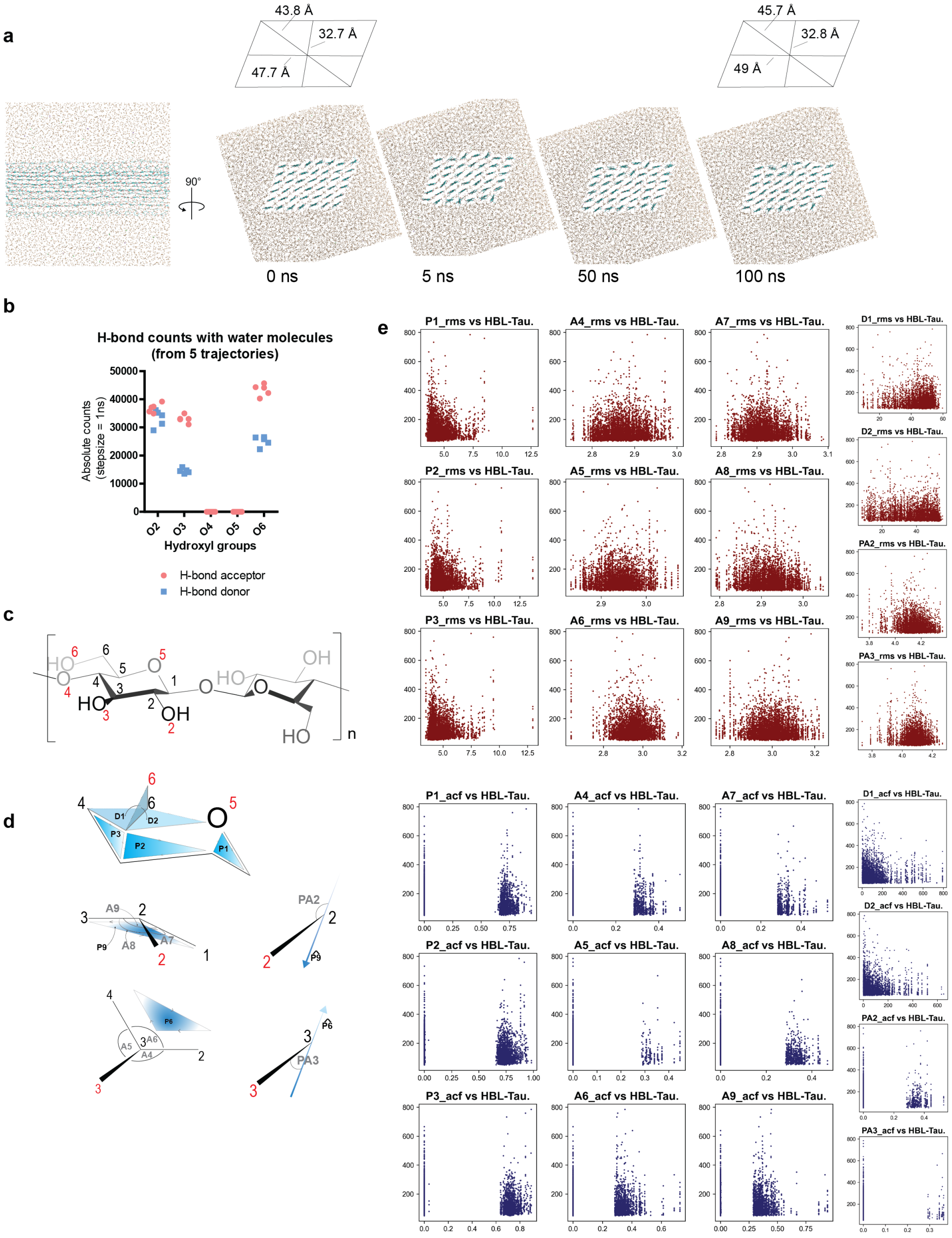

**Extended Data Fig. 4| MD simulation setup of a 42-chain fibril containing 18 glucosyl units per strand.** (**a**) Slight conformational rearrangements during the first few nanoseconds of unrestrained equilibration. (**b**) H-bonds between water and glucans were identified throughout 5 trajectories and plotted as frequency vs the glucan hydroxyls (labelled with oxygen annotation for glucopyranose) involved in forming them. Right panel: Snapshot after 500 ns simulation showing the coordination of three water molecules. (**c**) Numeric annotation of carbon (in black) and oxygen (in red) atoms. (**d**) Graphical description of the features in sugar moieties tracked in a trajectory. Prefix description: P= angular displacement of plane vector between two frames, A= absolute angle in a frame, PA = angle between ring carbon-hydroxyl and plane vector (shown as unit vector), D = dihedral angle. Numeric labels for endpoints follow atomic annotations from (c), feature suffixes were randomly assigned. (**e**) Scatter plots of longest H-bond lifetimes (HBL-Tau, Y axes) vs the features defined in (d) (X axes) of the sugar unit involved in the H-bonding. Suffix description: _rms = root mean squared fluctuation of the feature (Prefix), _acf = autocorrelation function constant of the feature. Acf values are in picoseconds (ps), rms values are in radians. While a distinct linear correlation is not seen between HBL-Tau and any feature, the variation in rms as well as acf taper with increasing HBL-Tau, suggesting that the participating sugars converge to uniform thermal fluctuations within an ordered domain.

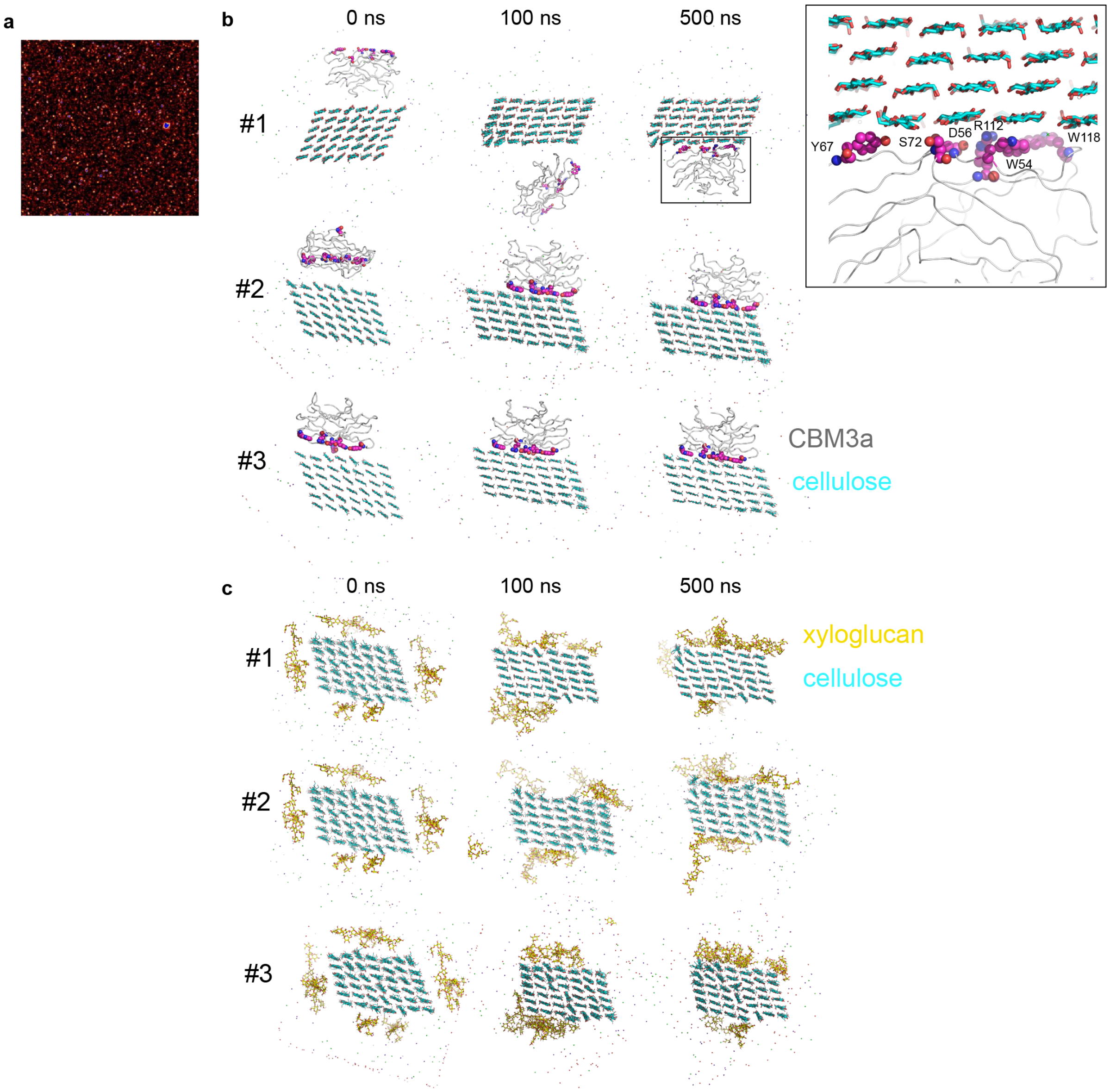


**Extended Data Fig. 5| CBM3a and xyloglucan binding to the tunicate cellulose fiber.** (**a**) A control experiment of tamarind xyloglucan incubated with CBM3a-GFP does not reveal fibrous structures. (**b**) CBM3a (PDB 4JO5) was placed in three different orientations into the simulation box together with a 42-chain hydrated cellulose fibril, followed by 500 ns of unrestraint simulation. (**c**) Three repeats of an unrestraint 500 ns simulation of the cellulose fibril in the presence of six xyloglucan XXXGXXX oligosaccharides with structure [Xyl-α-(1,6)]-Glc-β-(1,4)-[Xyl-α-(1,6)]-Glc-β-(1,4)-[Xyl-α-(1,6)]-Glc-β-(1,4)- Glc-β-(1,4)-[Xyl-α-(1,6)]-Glc-β-(1,4)-[Xyl-α-(1,6)]-Glc-β-(1,4)-[Xyl-α-(1,6)]-Glc-β (shown as yellow sticks).

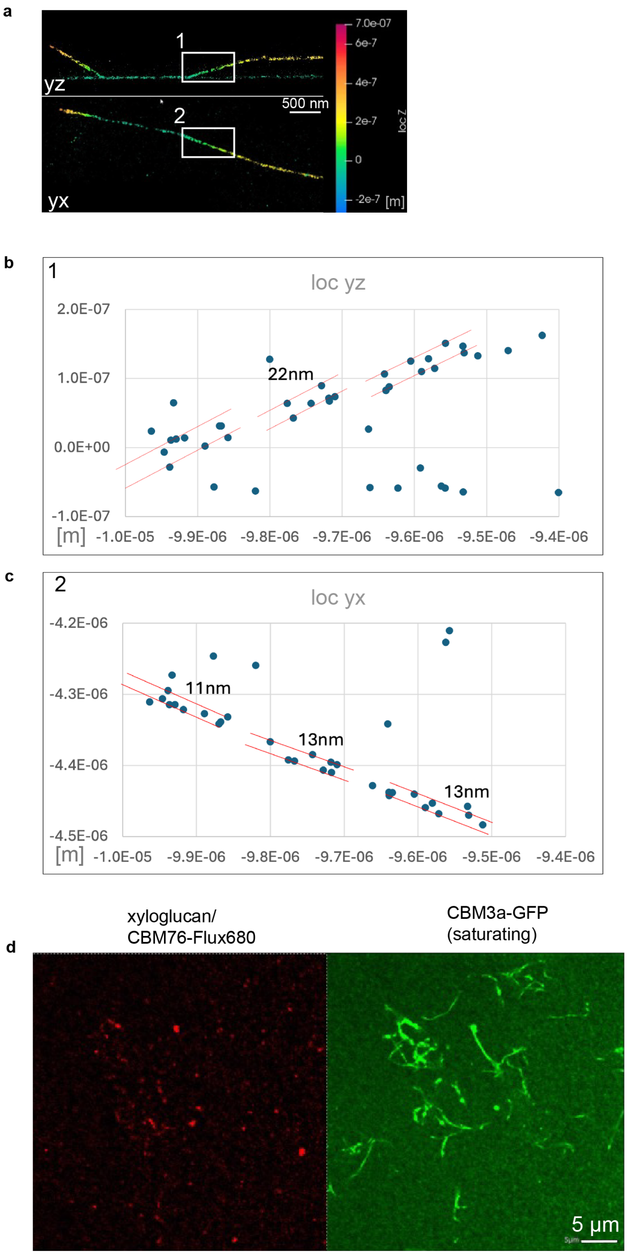


**Extended Data Fig. 6| Minflux nanoscopy of CBM3a-labeled cellulose fibers**. (**a**) Representative MINFLUX image of CBM3a-SNAP-AF647 labeled fibers with the regions used for analyses boxed in white. (**b**, **c**) Localizations of the position-averaged fluorophores identified in regions 1 and 2. (**d**) Confocal images of cellulose fibers first incubated with saturating CBM3a-GFP and then with xyloglucan/CBM76-Flux 660. CBM3a substantially reduces xyloglucan binding.

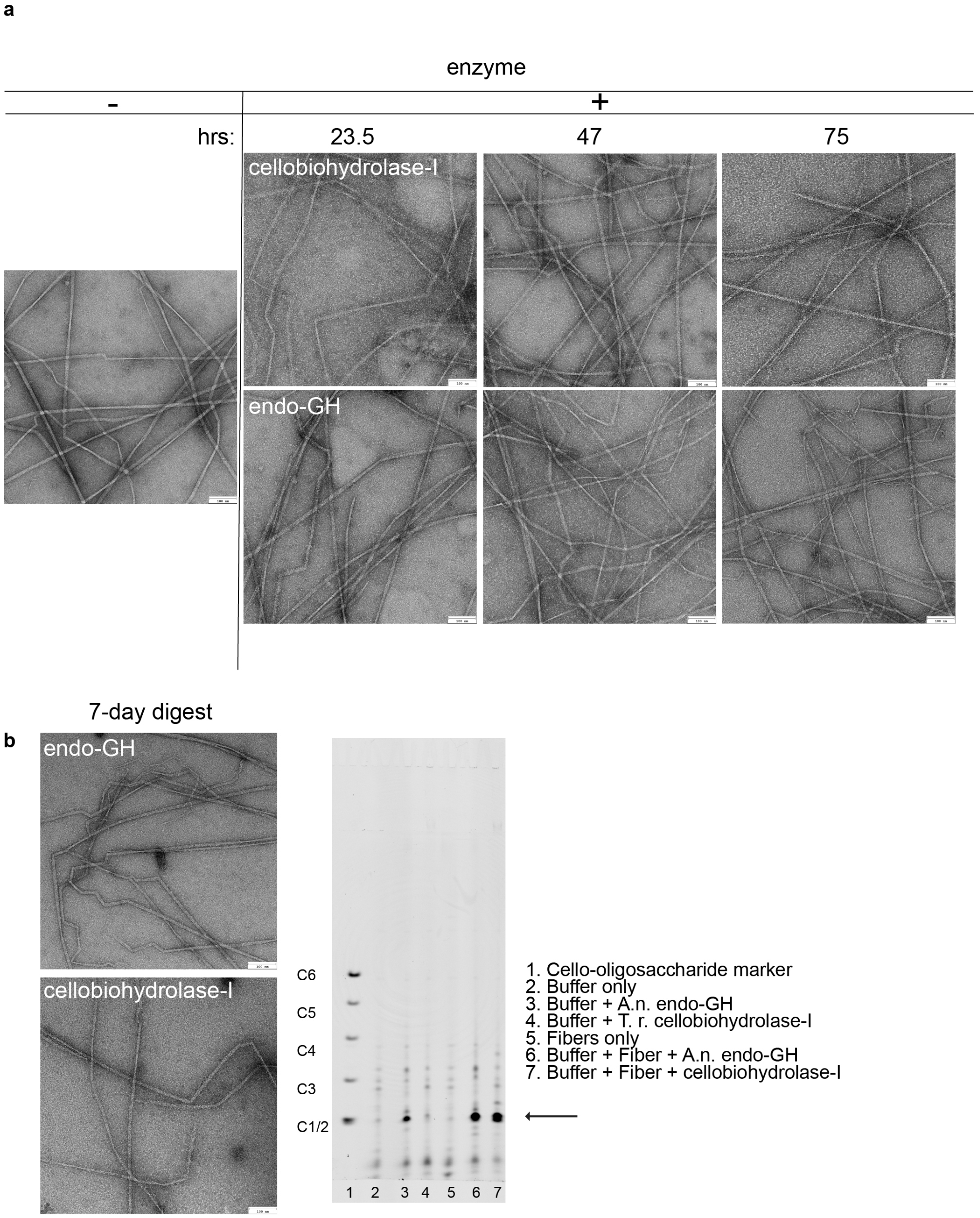


**Extended Data Fig. 7| Enzymatic degradation of the hydrated tunicate cellulose fibers.** (**a**) Negative stain images of fibers incubated with the indicated enzymes for 23.5 to 75 hours. Cellobiohydrolase-I was from *Trichoderma reesei* and the endo-β-(1,4) glucanase (endo-GH) was from *Aspergillus niger*. (**b**) Polysaccharide analysis by carbohydrate electrophoresis (PACE) of cellulase treated tunicate fibers. Left: Negative stain images of the tunicate fibers after incubation with the indicated enzymes for seven days. Right: PACE analysis of the same sample after 7 days indicating the release of mono- or disaccharide units.

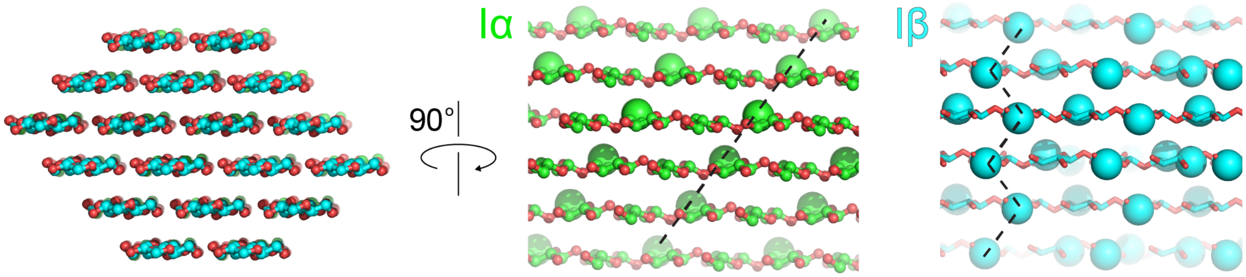


**Extended Data Fig. 8| Comparison of cellulose $I_\alpha$ and $I_\beta$.** The corresponding coordinates were obtained from the Cambridge Small Molecule Database (deposition numbers 792796 and 810597) and symmetry expanded in PyMOL. Segments corresponding to an 18-chain microfibril in the 2-3-4-4-3-2 configuration were selected and both allomorphs were aligned in PyMOL (left panel). Allomorphs are colored green and cyan for the carbon atoms of $I_\alpha$ and $I_\beta$, respectively. The side views show the glucosyl units' C6 carbons as spheres, for orientation. Dashed lines indicate the shifts of the glucan strands in adjacent layers.

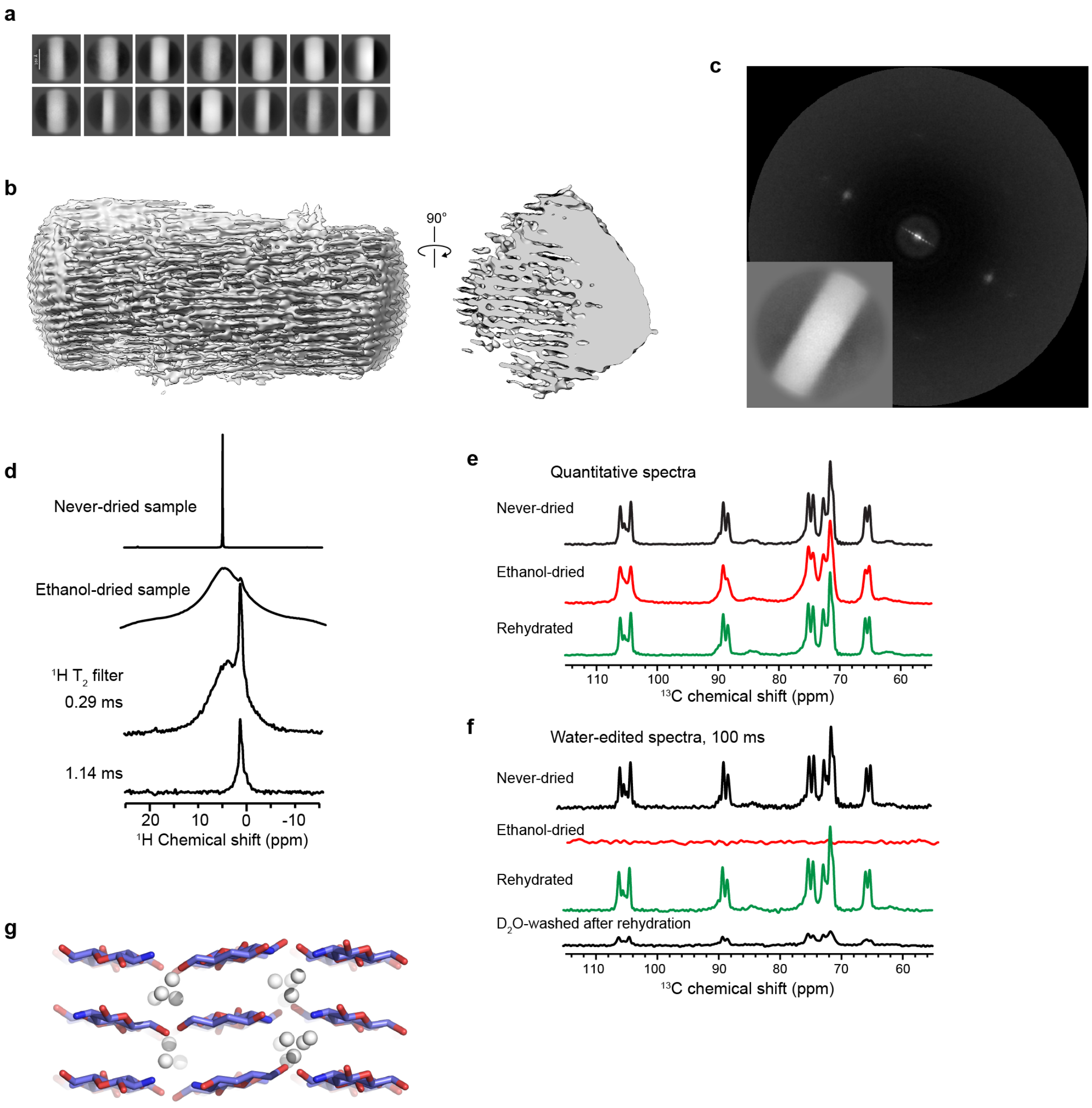


**Extended Data Fig. 9| Analysis of dried tunicate cellulose fibers.** (**a**) 2D class averages of dried tunicate fibers. (**b**) 3D reconstruction of a dried tunicate fiber. (**c**) Power spectrum of the indicated 2D class average calculated in cryoSPARC. (**d-f**) ssNMR analysis of the dried and rehydrated cellulose fibers. (**d**) $^{1}H$ NMR spectra of ethanol-dried sample with varying $^{1}H$ $T_2$ filter times compared to the never-dried spectrum. The dried sample no longer has a narrow water peak, confirming the quantitative removal of bulk water. The remaining broad $^{1}H$ peak, centered at 4.37 ppm, can be attributed to cellulose protons. Application of a $^{1}H$ $T_2$ filter with increasing duration progressively suppressed the intensity of the broad $^{1}H$ peak, exposing a sharp peak at 1.23 ppm, which may reflect the cellulose hydroxymethyl protons. (**e**) $^{13}C$ spectra of never dried, ethanol-dried and rehydrated tunicate cellulose. All three spectra show identical $^{13}C$ chemical shifts, but the dried sample has broader linewidths, indicating conformational disorder. Rehydration restored the linewidths of the never-dried sample, indicating that rehydration restored the conformational homogeneity of the original never-dried cellulose. The three spectra are plotted to match the intensity of the 71.6-ppm peak. (**f**) Water-edited $^{13}C$ spectra after 100 ms $^{1}H$ mixing for four differently treated tunicate cellulose samples. Ethanol drying suppressed the dynamic water $^{1}H$ magnetization, thus giving no $^{13}C$ signals. Rehydrated sample reached similar water-transferred intensities as the never-dried sample, indicating similar water accessibility of the fibrils. $D_2O$ washing after rehydration retained about 20% of the intensities of the rehydrated intensities, indicating that about ~20% of the water resides within the fibril interior, similar to the estimated 30% of the water content in the native never-dried tunicate cellulose. (**g**) Structure of hydrated chitosan with water molecules shown as gray spheres, from reference [43].

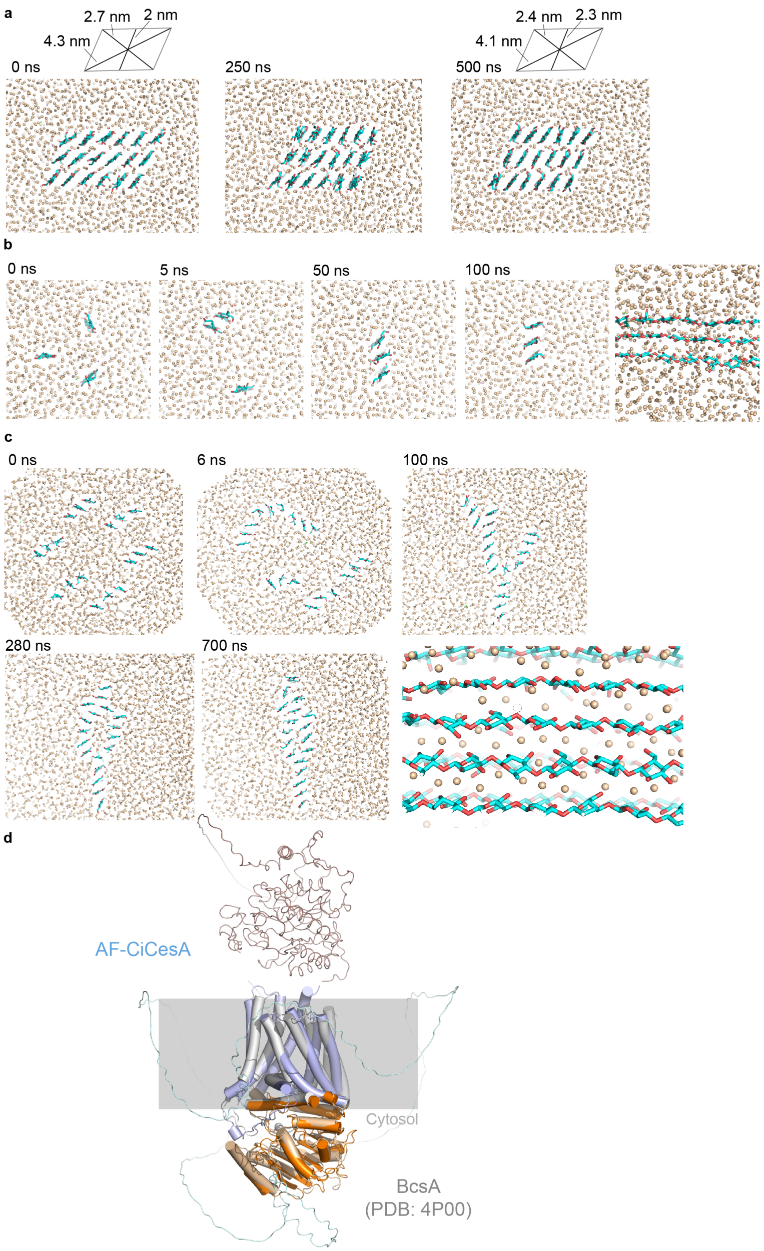


**Extended Data Fig. 10| Cellulose fibrillogenesis.** (**a**) MD simulation of an 18-chain hydrated cellulose fibril. The fibril is stable after an initial subtle rearrangement. (**b**) MD simulation of three isolated glucan strands under periodic boundary conditions. The formed protofibril is stable for > 500 ns. (**c**) MD simulation of six protofibrils (containing three glucans each) for 500 ns. Four out of eight simulations show lateral associations of cellulose strands with water molecules at the interface (shown in the side view of the 700 ns snapshot). (**d**) AlphaFold3 predicted structure of Ciona cellulose synthase overlayed with *Rhodobacter sphaeroides* BcsA (PDB: 4P00). BcsA is colored gray and pale orange, and Ciona cellulose synthase is shown in orange and blue for its catalytic domain and transmembrane region, respectively. The tunicate enzyme contains an N-terminal extension and a C-terminal extracellular cellulase domain (colored brown).
